# Mechanistic basis for the opposing effects of H2A and H2B ubiquitination on nucleosome stability and dynamics

**DOI:** 10.1101/2025.02.13.638112

**Authors:** Lokesh Baweja, Jeff Wereszczynski

## Abstract

Nucleosome ubiquitination at lysine 119 of histone H2A (H2AK119ub) and lysine 120 of histone H2B (H2BK120ub) are prominent post-translational modifications with opposing roles in chromatin regulation. Although H2AK119ub is associated with transcriptional repression and H2BK120ub with activation, the molecular basis for these contrasting effects has remained unclear. Here, we use microsecond all-atom and millisecond coarse-grained molecular dynamics simulations to reveal how the position of ubiquitin reshapes nucleosome structure and assembly. H2AK119ub rigidifies the histone core by indirectly reinforcing the L1–L1 interface between H2A histones, strengthening both tetramer–dimer and dimer–dimer interactions, and slowing complete nucleosome assembly. In contrast, H2BK120ub disrupts these interfaces, weakens the histone core, and favors partially assembled hexasome and tetrasome states. Both modifications cause dramatic slowdowns in nucleosome folding, with H2BK120ub producing an order-of-magnitude greater effect. These simulations establish clear molecular mechanisms by which site-specific ubiquitination alters nucleosome stability and assembly kinetics. Our findings quantitatively explain how H2A and H2B ubiquitination exert opposing effects on chromatin regulation. This mechanism is directly relevant to the opposing roles of these marks in transcriptional activation and repression, and may represent one way that combinations of histone modifications modulate chromatin function *in vivo*.

## Introduction

The nucleosome core particle (NCP) is the fundamental unit of chromatin, where approximately 147 base pairs of DNA are tightly wrapped around a histone core. This core is formed by the association of an (H3/H4)_2_ tetramer with two H2A/H2B dimers^1–4^. Each histone consists of three alpha helices (a1, a2, and a3) and disordered histone “tails” that protrude from the core^5^. At physiological salt concentrations, the histone octamer is only stable when DNA is wrapped around it due to the high positive charges of the histones^6,7^.

The strong association of DNA with the histone core blocks DNA-binding sites required for vital cellular processes like transcriptional regulation and DNA repair^8–11^. One mechanism by which histone-DNA and histone-histone interactions are modulated to tune DNA accessibility is through histone post-translational modifications (PTMs)^12,13^. These modifications vary in both type and size; acetylation, methylation, and phosphorylation are small chemical modifications, while ubiquitination involves the addition of 76 amino acids through an isopeptide bond and introduces both steric bulk and new residues for potential interactions within the nucleosome^14–16^. Ubiquitination occurs predominantly on H2A and H2B, accounting for approximately 10-15% of the total histone PTMs^17–19^, while less than 0.3% and 0.1% of the histones H3 and H4^20^, respectively, are ubiquitinated. Two major histone ubiquitination sites have been identified in higher organisms: lysine 119 on H2A (H2AK119ub) and lysine 120 on H2B (H2BK120ub)^21,22^. H2A119Kub is located on the disordered C-terminal tail of H2A near the dyad axis and the entry and exit DNA, whereas H2BK120 is on an H2B alpha helix near the acidic patch (Figure 1).

**Figure 1:**
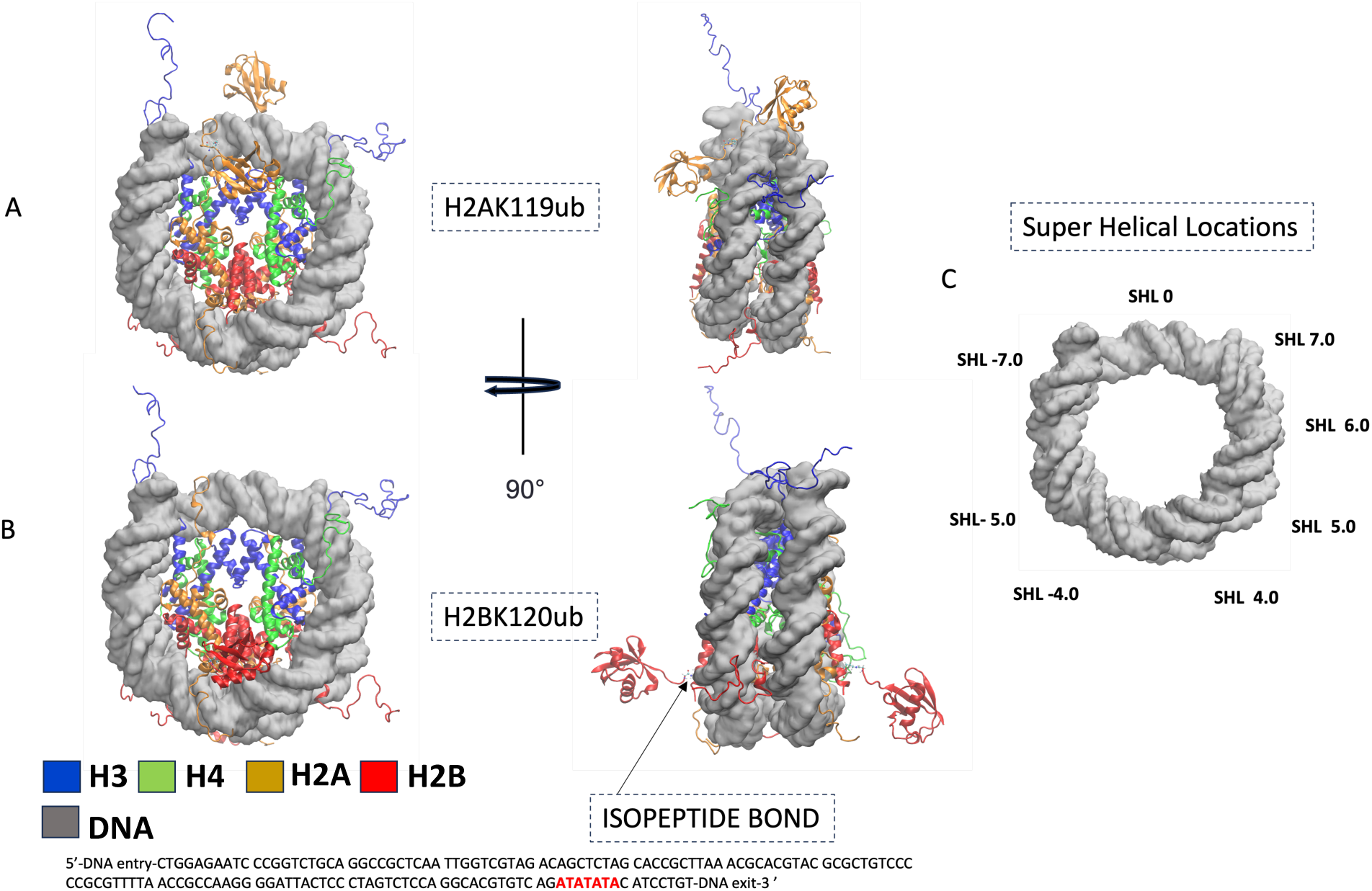
(A-B) Initial conformations of the H2AK119ub and H2BK120ub nucleosome systems generated with Chimera and the Widom 601 nucleotide sequence in 5’ to 3’ orientation. The DNA exit end is AT-rich. (C) Super helical locations (SHLs) are sites marked on the nucleosomal DNA.

Recent chromatin immunoprecipitation sequencing by Tamburri *et al.* demonstrated that H2AK119 ubiquitination co-exists with H3K27 trimethylation in transcriptionally inactive heterochromatin regions^23^. H2AK119 is also required for high-affinity binding of the RING1 and YY1 binding protein (RYBP) of the Polycomb repressive complex, which is implicated in heterochromatin formation^24^. Single-molecule FRET experiments by Xia *et al.* showed that H2AK119ub reinforces the mechanical stability of the nucleosome, although only weak interactions of ubiquitin with DNA were reported^25^. In contrast, H2BK120 ubiquitination is associated with actively transcribed regions and chromatin decompaction^26^. H2BK120ub promotes the binding of the histone chaperone FACT (Facilitates Chromatin Transcription) complex by decreasing nucleosomal stability^26^.

Despite these excellent experimental advances, the mechanisms by which ubiquitination modulates nucleosome dynamics across different timescales remain unclear^27,28^. Recent all-atom and coarse-grain molecular dynamics (MD) simulations have begun to detail the effects of histone modifications on nucleosome behavior and gene regulation^29–36^. However, the physical mechanisms by which ubiquitination, notably H2AK119ub and H2BK120ub, modulate nucleosome stability and dynamics are still not well understood.

Here, we dissect the site-specific mechanisms by which H2AK119ub and H2BK120ub alter nucleosome structure, stability, and folding, using microsecond all-atom and millisecond coarse-grained MD simulations combined with Markov state modeling (MSM)^37^. While our simulations cannot capture every aspect of the cellular environment, they allow us to directly quantify the effects of these modifications at resolutions inaccessible to experiment. Our all-atom simulations show that H2AK119ub indirectly stabilizes the nucleosome by increasing contacts at the L1–L1 interface between H2A histones and reinforcing both tetramer–dimer and dimer–dimer interactions, while H2BK120ub disrupts these interactions and destabilizes the core. Coarse-grained simulations further demonstrate that H2BK120ub shifts the equilibrium toward partially assembled hexasomal and tetrasomal intermediates and that both modifications dramatically slow nucleosome folding. These results quantitatively link specific histone ubiquitination events to the energetic landscape of nucleosome assembly, providing a mechanistic basis for the antagonistic regulation of chromatin by H2A and H2B ubiquitination.

## Materials and Methods

### System preparation and simulations setup

Initial structures of the canonical and H2AK119 and H2BK120 ubiquitinated nucleosome core particle (NCPs) were constructed using Chimera based on the 1KX5 NCP structure,^38,39^ where the DNA was substituted with a Widom 601 DNA sequence.^40^ H2AK119ub and H2BK120ub systems were generated using Chimera by conjugating the ubiquitin moieties (as resolved in PDB ID 3H7S) at the sites H2AK119 and H2BK120 with an isopeptide linkage.^39^ Histone tail conformations from the 1KX5 structure were used. Missing residues were modeled, and protonation states were assigned using tleap from the AMBER suite^41^. The parameters for the isopepetide linkage were obtained with antechamber using GAFF with partial charges derived from restrained electrostatic potential fitting at the HF/6-31G* using gaussian level 09^42^.

The protein and DNA in these systems were modeled with the Amber ff19SB^43^ and BSC1^44^ force fields, respectively. All systems were then solvated with OPC water models^45^ with 0.15M KCl. A 4fs timestep was used in conjunction with hydrogen mass repartitioning.^46,47^ All systems were first energy minimized for 5000 steps with solute harmonic restraints of 10 kcal/mol/A, followed by 5000 steps with no restraints. Minimized systems were then equilibrated for 100 ps by keeping the volume constant and gradually increasing the temperature from 10K to 310K with heavy atom restraints. Afterwards, the heavy atom restraints were gradually decreased over 1000 ps in the isothermal and isobaric ensemble. Production runs initiated from independent randomized velocities were performed in triplicate for 2 µs with Amber 22, accumulation 18 µs of sampling across all three systems (Table S1). Trajectories were recorded every 10 ps and visualized using VMD^48^ and PyMol.^49^ The last 1.5 µs were analyzed using CPPTRAJ^50^ and Python scripts allowing for 500 ns of equilibration.

Coarse grain (CG) simulations used the AICG2+ protein potential^51^ and 3SPN2.C DNA sequence-dependent potential models.^52^ In these models, each amino acid is treated with a single Ca bead and each nucleotide consists of three beads corresponding to the base, sugar, and phosphate. The flexible histone tails (residues 1-32 for H3, 1-23 for H4, 1-14 and 121-128 for H2A, 1-26 for H2B) were treated with the flexible local potential. The DNA and histones interacted via excluded volume, Debye–Hückel electrostatics and hydrogen bond interactions.^53^ The charges for the structured histone core were derived using the RESPAC method,^54^ whereas for the flexible tails and ubiquitin moiety we used standard integer charges. Scaling factors of 0.3 and 0.6 were used for non-bonded interactions between the (H3/H4)_2_ tetramer and H2A/H2B dimers and the DNA/histones, respectively, as used in previous work.^31^

For coarse-graining of ubiquitinated systems, the isopeptide linkage was treated using the flexible local potential present in the cafemol package. The parameter for this linkage were derived from all-atom simulations of K63 linked diubiquitin (PDB ID:3H7S).^55^ We coarse-grained the isopeptide bond by placing a Ca bead at the center of mass of the –GLY-LYS– linkage (Figure S2). We then used a protocol adopted from Terakawa and Takada based on Boltzmann Inversion to obtain (GLY-ISO-LYS) angle and (–GLY-GLY-ISO-LYS–) and (–GLY-ISO-LYS-LYS–) dihedral parameters.^56^ We carried out five Boltzmann inversion simulations to capture the bond angle and dihedral distributions obtained from all-atom simulations (Figure S2).

The initial tetrasomal (T), hexasome-left (HL), hexasome-right (HR), and Nucleosome (N) conformations for CG simulation were obtained from our previous study (Figure S1).^57^ CG molecular dynamics simulations were conducted using a modified version of CafeMolv3.1 software,^58^ to treat the -ISO-bead with flexible local potential. Integration of the equations of motion was performed through Langevin dynamics at a temperature of 310 K, employing a timestep of 0.3 CafeMol time units. All units in coarse-grained simulations are reduced units, where the time step qualitatively corresponds to approximately 30 ps.^31^ During CG simulations, harmonic restraints were applied to the (H3/H4)_2_ tetramer and dyad base pair to ensure the stability of the tetrasomal system. Specifically, the structured region of the two H3 histones experienced restraints with a force constant of 0.001 kcal/mol/Å^2^.^31^ Additionally, harmonic restraints were imposed on the phosphate beads of the DNA dyad residues with a force constant of 0.1 kcal/mol/Å^2^. These restraints limited large-scale sliding of DNA relative to the (H3/H4)_2_ tetramer while leaving the unbound H2A/H2B dimers unrestricted. This setup is used to mimic the experimental conditions used for nucleosome refolding studies from the tetrasomal states.^59^

### Markov State Modeling

We modelled nucleosome reassembly from tetrasomal states using a set of six collective variables that defined the binding of H2A/H2B dimers with the (H3-H4)_2_ interface and the degree of DNA opening. The first two collective variables measured the normalized number of contacts for each dimer with the tetramer. To do this, we measured the distance between the center of mass of the H2B residues 73-100 and the center of mass of the H4 residues 65-92 and normalized it to between 0 and 1. A value close to 1 indicates a dimer bound to the (H3/H4)_2_ tetramer in its native configuration and a value close to 0 indicates no dimer/tetramer interactions. The third and fourth collective variables represent the radius of gyration of the entry and exit halves of the DNA. The entry half includes base pairs from the DNA end up to the dyad, while the exit half consists of the remaining segment beyond the dyad. We also defined two other collective variables based on the distance between the DNA entry and the DNA exit ends from the dyad. Although the two H3/H4 binding interfaces were theoretically identical, the asymmetry introduced by the difference in the composition of 601 DNA wrapped around the tetramer led us to distinguish between the DNA entry and AT-rich DNA exit sides of the nucleosome.

We built Markov State Models (MSMs) for nucleosome assembly at a salt concentration of 300 mM, which was lower than the 400 mM concentration used in a previous study on nucleosome assembly, due to the dramatic destabilization of our H2BK120ub system at 400 mM. We aggregated all 800 CG MD trajectories, followed by dimensionality reduction using time-lagged independent component analysis (TICA)^60^ with a lag time of 2 *↑* 10^6^ as implemented in PyEMMA 2,^61^ and clustered the conformations into 400 discrete states using k-means.^62^ For all systems, the MSMs were generated using a lag-time of 2 *↑* 10^6^ time-steps. Further increases in the lag-time did not lead to significant changes in the MSM implied time scales (Figure S4). However, the H2AK119ub and H2BK120ub systems led to faster convergence due to the higher transitions of the nucleosome to hexasomal states. Furthermore, the first two TICA coordinates were sufficient to separate the system into four regions representing the T, HR, HL, and N nucleosome states (Figure S3).

### Simulation Analyses

Root mean square deviations (RMSDs) of the ubiquitin Ca atoms were calculated using CPPTRAJ^50^ after aligning the trajectories to the (H3-H4)_2_ tetramer excluding tails. Dimer-dimer and dimer-tetramer contact analyses were conducted using the MDAnalysis package^63^ and visualized using matplotlib^64^, where contacts were defined between histone heavy atoms that were within 4.5 Å of one another. The MM-GBSA (Molecular mechanics generalized born surface area) analyses were performed with igb=5 and a salt concentration of 0.15 M with the MMPBSA.py program^65^. Error bars represent the standard error of the mean, with a decorrelation time of 12.5 ns calculated by a statistical inefficiency test^66^. Generalized correlation analysis of DNA and histones were performed on C1’ and Ca, respectively using correlation plus^67^. Difference maps of H2A119Kub and H2BK120ub systems were generated after subtracting from the canonical maps. Coarse-grained simulations were analyzed with in-house generated python scripts.

During manuscript preparation, the authors used ChatGPT to assist with language editing, text refinement, and readability improvements. All content was reviewed and edited by the authors, who take full responsibility for the final text.

## Results

### H2AK119ub Exhibits Stronger DNA Interactions than H2BK120ub

We conducted a series of simulations to investigate the mechanisms by which H2AK119ub and H2BK120ub affect nucleosome structures and dynamics. Specifically, we ran three replicates of 2 µs each using all-atom simulations for the canonical nucleosome core particle (NCP) and for NCPs ubiquitinated at H2AK119 (H2AK119ub) and H2BK120 (H2BK120ub) sites. In both modified systems, we observed considerable movement of the ubiquitin moieties. Root mean square deviation (RMSD) measurements of each ubiquitin relative to the core showed displacements ranging from 15 to 50 Å in the H2AK119ub system and from 10 to 30 Å in the H2BK120ub system (Figure 2).

**Figure 2:**
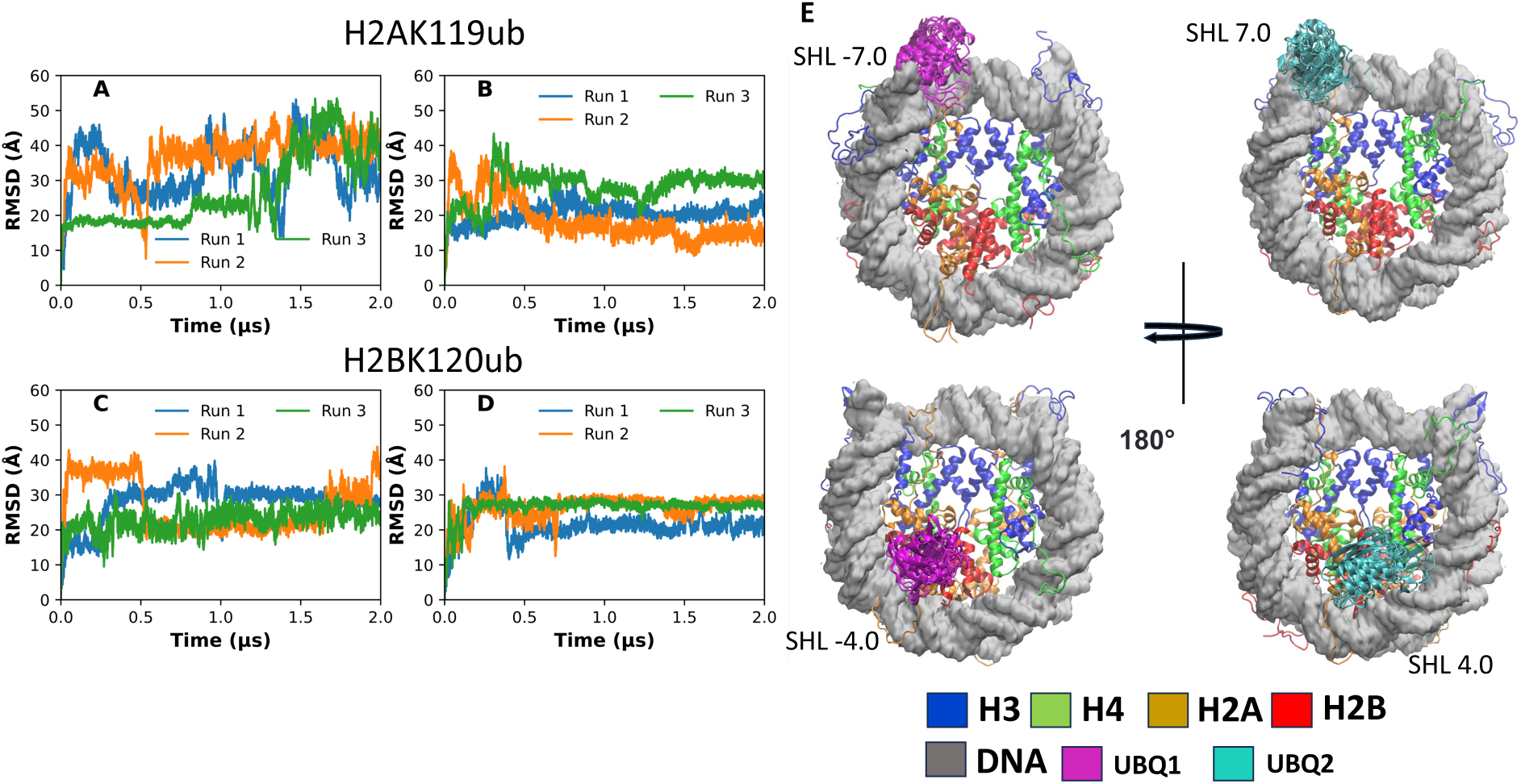
(A–D) Root mean squared deviations (RMSD) of ubiquitin in H2A119ub and H2BK120ub nucleosome systems, which quantify its motion relative to the histone core. Ubiquitin has more restricted dynamics in the H2BK120ub system. (E) Representative configurations from the five most populated structural clusters, based on RMSD in the equilibrated portions of the trajectories.

Analysis of ubiquitin/DNA interactions revealed different patterns based on the ubiquitination site. In the H2AK119ub system, ubiquitins interacted primarily with the entry DNA and the AT-rich exit DNA end. The entry-side ubiquitin bound to superhelical locations (SHL) −7.0, while the exit-side ubiquitin bound to SHL 7.0 (Figure S5). Both also made contacts near the DNA dyad at SHLs 0.0 and −0.5. In contrast, the H2BK120ub system showed fewer DNA contacts and differences in the SHL interaction pattern. The exit-side ubiquitin did not make significant contact with DNA, while the entry-side ubiquitin interacted with SHLs 3.5 and 4.0.

To assess the strength of these binding interactions, we calculated MM/GBSA-derived interaction energies between each ubiquitin and the nucleosomal DNA and histone core (Table 1). In the H2AK119ub system, the entry side ubiquitin had a favorable interaction of −27.1*±*0.4 kcal/mol with DNA and an unfavorable interaction of 8.2*±*1.1 kcal/mol with the histone core, for a total interaction of −15.9*±*1.1 kcal/mol. The exit-side ubiquitin had a similarly favorable interaction energy of −19.4*±*kcal/mol with DNA and an unfavorable interaction of 11.1*±*1.1 kcal/mol with the histone core, for a total of −8.3*±*1.1 kcal/mol. Conversely, in the H2BK120ub system, interaction energies were significantly less favorable. The exit-side ubiquitin had a DNA interaction of −3.3*±*0.3 kcal/mol and histone core interaction of 4.6*±*0.8 kcal/mol, for a total of 3.4*±*0.3 kcal/mol, while the entry-side ubiquitin had unfavorable interactions with both the DNA and histone core of 7.5*±*0.6 and 6.2*±*1.0 kcal/mol, repectively, for a total interaction energy of 13.5*±* 1.0 kcal/mol.

**Table 1:** MM/GBSA-derived binding energies (kcal/mol) for histone–DNA and histone–histone interfaces in each nucleosome system, averaged over three independent simulations. Energy terms include van der Waals (ΔE_vdW_), electrostatic (ΔE_ele_), and total interaction energy (ΔE_total_).

| System | Complex | $\Delta E_{\text{vdW}}$ | $\Delta E_{\text{ele}}$ | $\Delta E_{\text{total}}$ |
| --- | --- | --- | --- | --- |
| H2AK119ub | DNA-ubiquitin1 | -37.5±0.1 | 10.4±0.4 | -27.1±0.4 |
| | DNA-ubiquitin2 | -41.8±0.1 | 22.4±0.3 | -19.4±0.3 |
| | core-ubiquitin1 | -15.8±0.1 | 24.0±1.1 | 8.2±1.1 |
| | core-ubiquitin2 | -26.5±0.1 | 37.5±1.0 | 11.0±1.1 |
| | dimer1-hexamer | -221.4±0.2 | 211.3±1.1 | -10.1±1.1 |
| | dimer2-hexamer | -222.4±0.2 | 207.0±0.8 | -15.4±0.8 |
| | dimer-dimer | -39.4±0.1 | 44.0±0.9 | 4.6±0.9 |
| H2BK120ub | DNA-ubiquitin1 | -10.0±0.1 | 6.7±0.3 | -3.3±0.3 |
| | DNA-ubiquitin2 | -2.0±0.1 | 9.5±0.6 | 7.5±0.6 |
| | core-ubiquitin1 | -65.4±0.2 | 70.0±0.8 | 4.6±0.8 |
| | core-ubiquitin2 | -46.0±0.1 | 52.0±1.0 | 6.0 ±1.0 |
| | dimer1-hexamer | -202.7±0.1 | 206.4±0.8 | 3.7±0.8 |
| | dimer2-hexamer | -202.0±0.1 | 203.3±0.8 | 1.3±0.8 |
| | dimer-dimer | -22.0±0.1 | 36.1±0.8 | 14.1±0.8 |
| Canonical | dimer1-hexamer | -210.7±0.1 | 206.0±0.5 | -4.7±0.5 |
| | dimer2-hexamer | -208.7±0.1 | 210.0±0.6 | 1.3±0.6 |
| | dimer-dimer | -28.2±0.1 | 41.5±0.8 | 13.2±0.8 |

**Table 2:** Relative free energies (kcal/mol) of principal macrostates along the canonical nucleosome folding pathway, calculated from Markov State Model analysis. Energies are given relative to the most stable state.

| State | Kcal/mol |
| --- | --- |
| Tetrasome (extended), | T (1.29), T2 (1.40) |
| Tetrasome (Compact) | T1 (1.57) |
| Hexasome Right (HR) - DNA-exit-side folded | HR (0.25), HR2 (3.04), HR3 (3.06) |
| Hexasome Left (HL) - DNA-entry-side folded | HL (2.18) |
| Nucleosome (N) | N (0.00) |

A decomposition of these energies into van der Waals and electrostatic components revealed that van der Waals forces were the dominant contributors to ubiquitin-DNA binding, while electrostatic contributions were consistently unfavorable (Table 1). Although MM/GBSA calculations involve several approximations, the qualitative nature of these results is consistent with previous experimental studies, which show weak binding of ubiquitin to nucleosomal DNA.^68^ Together, these findings suggest that ubiquitins on the C-terminal H2A tails sample more DNA binding sites and form stronger interactions than those bound to H2BK120.

### Distinct Ubiquitination Sites Drive Opposing Dimer Motions

Having established that ubiquitin moieties undergo significant motions on the µs timescale, we next examined how ubiquitination affects the dynamics of the nucleosome core. To assess its impact on the H2A/H2B dimers, we quantified their motions relative to the tetramer by defining an angle between the vectors formed by the a2 helix of H2A and the dyad axis. This angle reflects inward or outward tilting of the dimers, with larger values corresponding to outward tilting and smaller values to motion toward the core (Figure 3).

**Figure 3:**
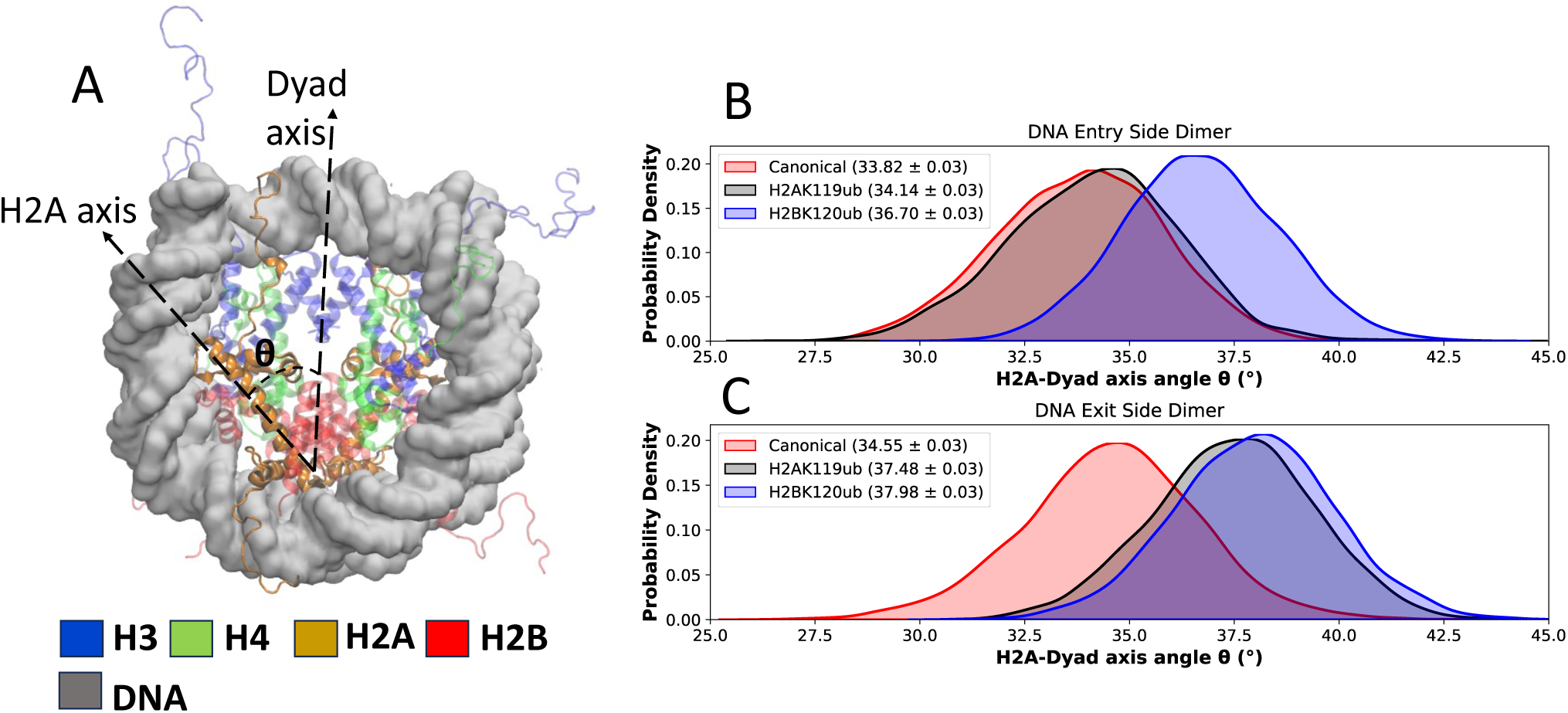
(A) Schematic of the dyad axis and the H2A axis, illustrating the angle measured to quantify dimer motion within the nucleosome core. (B–C) Distribution and mean values of the H2A-dyad angle. Both H2AK119ub and H2BK120ub systems exhibit increased outward tilting of the H2A–H2B dimers relative to the canonical nucleosome.

The mean tilting angles for the canonical system were 33.8*^→^* on the DNA entry side and 34.1*^→^* on the DNA exit side. In the H2AK119ub system, the entry-side dimer sampled values similar to the canonical system, but the exit-side dimer exhibited an increase of approximately 3*^→^*, suggesting an asymmetry in the dimer motions (Figure 3). In H2BK120ub, both dimers showed a statistically significant increase of 2-4*^→^* relative to the canonical case, with values of 36.7*^→^* and 38.0*^→^*, respectively. These results indicate greater outward motions of both dimers in H2BK120ub compared to the canonical and H2AK119ub nucleosomes. Additionally, the range of tilting angles in the canonical system spanned 18*^→^*, whereas in both H2AK119ub and H2BK120ub this range was narrowed to 12*^→^*, suggesting that ubiquitination may restrict overall dimer flexibility.

### Ubiquitination Reshapes Global Nucleosomal Dynamics

To further investigate the impact of system-dependent dimer motions in the histone core, we calculated the total number of contacts formed between the dimers using a 4.5 Å cutoff between all heavy atoms (Figure S6). In the H2AK119ub system, the number of dimer-dimer contacts increased significantly from 51 in the canonical system to approximately 77. In contrast, in H2BK120ub, the number of dimer/dimer contacts was substantially reduced to about 34.

To identify specific regions responsible for these differences, we calculated per-residue contact maps (Figure 4). In the canonical NCP, dimer-dimer interactions primarily involve the H2A L1 and H2B a2-helix (Figure 4b). The contact maps reveal a significant enrichment of contacts between the H2A L1-L1 loops and the L1-H2A-a1 interfaces in H2AK119ub (Figure 4c). In contrast, H2BK120ub displayed a substantial loss of contacts at the dimer-dimer interface (Figure 4d), likely due to increased outward movement and greater separation between dimers in this system.

**Figure 4:**
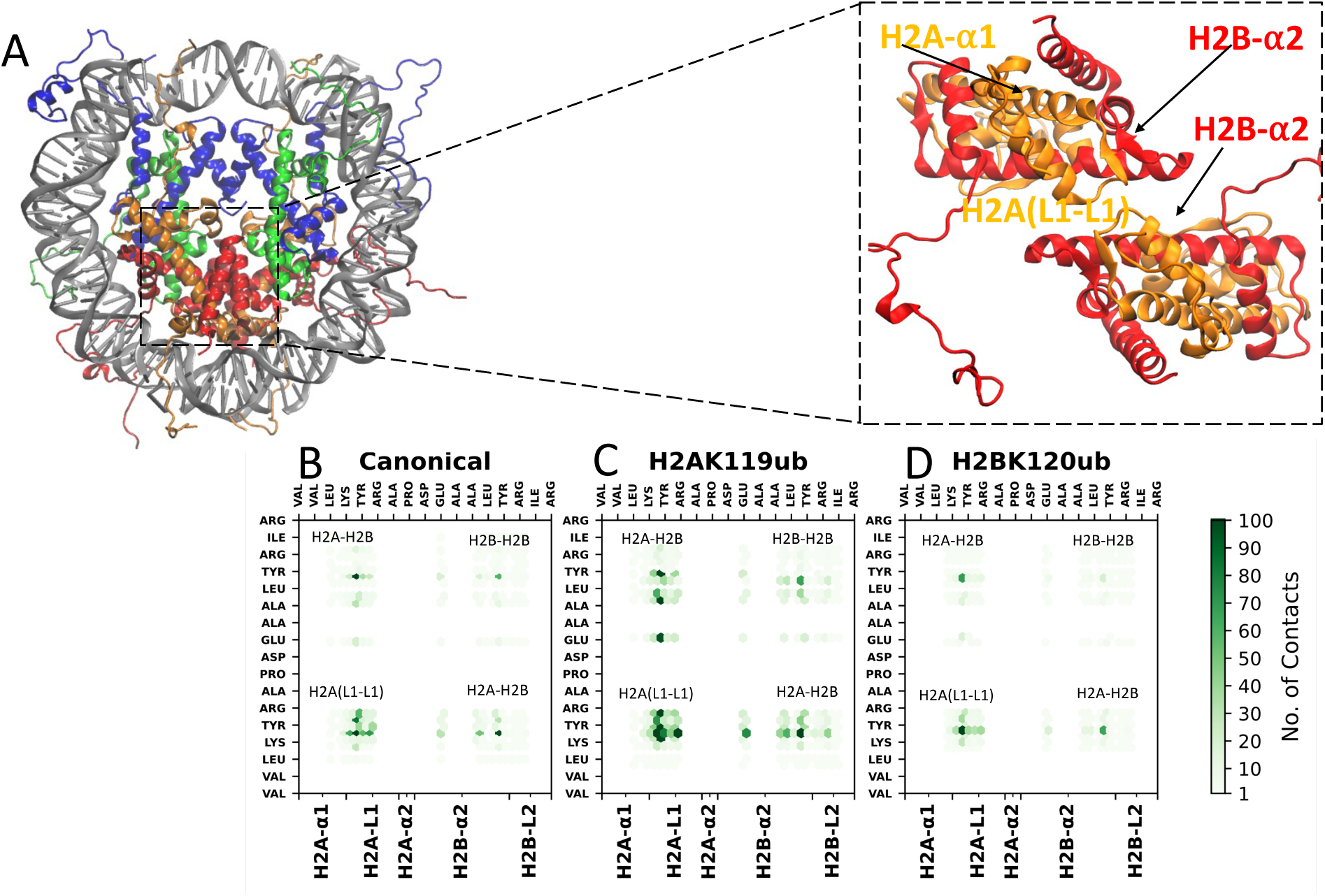
(A) Structural view of the H2A–H2B dimer–dimer interface in the nucleosome, highlighting the roles of the H2A L1–L1 loops and H2B a2 helices. (B–D) Per-residue contact maps showing inter-dimer interactions in the canonical, H2AK119ub, and H2BK120ub systems. The H2A L1–L1 region in H2AK119ub shows a pronounced increase in contacts. The color bar indicates the total number of contacts per residue pair.

To assess whether these motions also affect dimer-tetramer interactions, we analyzed the contacts at those interfaces. H2AK119ub showed a slight increase in dimer/tetramer contacts, from 431 in the canonical system to 440. In H2BK120ub, contacts decreased to 425 and 421, lower than both the canonical and H2AK119ub systems, suggesting that ubiquitination exerts opposite effects on these interfaces (Figure S7).

We next evaluated how these changes influence each dimer’s stability within the histone core using MM/GBSA calculations between each dimer and the remaining hexamer. These analyses revealed a significant increase in net van der Waals (vdW) interaction energy in H2AK119ub, resulting in more favorable total interaction energies than in the canonical NCP, with total ΔE values of −10.1*±*1.1 kcal/mol and −15.4*±*0.8 kcal/mol for the DNA entry and AT-rich DNA exit dimers, respectively. In contrast, MM/GBSA analysis for H2BK120ub indicated a net unfavorable dimer binding, with ΔE values of 3.7*±*0.8 kcal/mol and 1.3*±*0.8 kcal/mol, suggesting reduced dimer association stability in H2BK120ub.

To understand the effects of ubiquitination on DNA dynamics, we examined DNA base-pair unwrapping along the superhelical axis over our 2.0 µs simulations (Figure 5). In the canonical system, unwrapping of the entry DNA was observed in a single trajectory, where four to eight base pairs unwrapped after 1.0 µs, while the exit side showed only the transient unwrapping of up to four base pairs. In the H2BK120ub system, unwrapping of around four base pairs from the entry DNA and ten from the exit side occurred in the first trajectory, with little or no unwrapping in the others. In contrast, the H2AK119ub system showed only slight unwrapping in a single trajectory, briefly exceeding four base pairs, with no unwrapping in the other simulations. Together, these results suggest that H2AK119ub stabilizes DNA wrapping relative to the canonical and H2BK120ub systems, likely due to additional interactions introduced by ubiquitin at the entry and exit sites, which help maintain a more compact structure.

**Figure 5:**
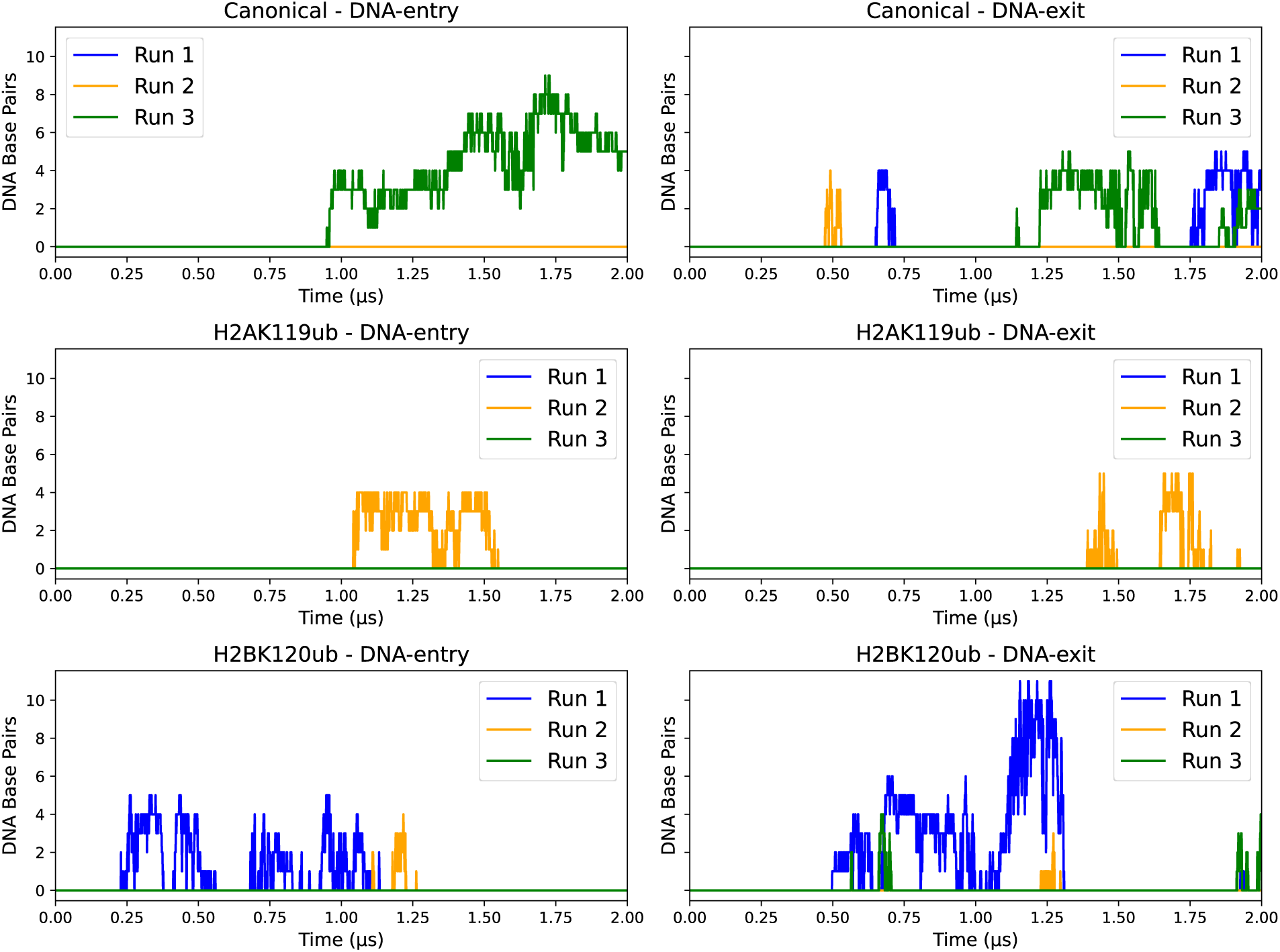
DNA unwrapping (“breathing”) at the nucleosome entry and exit sites, quantified by the number of base pairs unwrapped from the superhelical axis over time. Both the canonical and H2BK120ub nucleosomes show greater DNA unwrapping than the H2AK119ub system.

In addition to impacting dimer and DNA dynamics, ubiquitination also affected local and long-range dynamic correlations throughout the NCP (Figure S8). Mutual information calculations of protein Ca and DNA C1’ atom dynamics in the canonical system revealed high correlations between histone-histone and histone-DNA components. Inter-subunit correlations, represented by off-diagonal blocks in the correlation matrices, were lower for ubiquitinated states and drastically reduced in H2BK120ub.

Correlation differences relative to the canonical system show a disparate impact on system correlations from H2AK119ub and H2BK120ub ubiquitination. In H2AK119ub, the entry-side dimer had showed relatively small increases in correlation (on the order of 0.05-0.10) with the tetramer and DNA, while the exit dimer had lower correlations with the entire NCP (Figure 6a). In contrast, in H2BK120ub system, both exit and entry dimers showed a decoupling between H2A and H2B monomer dynamics, along with a greater reduction in the H2B correlations with the other histones and DNA (Figure 6b). At the same time, the N-termini of H3 and H4 had increased correlations with other histones and DNA.

**Figure 6:**
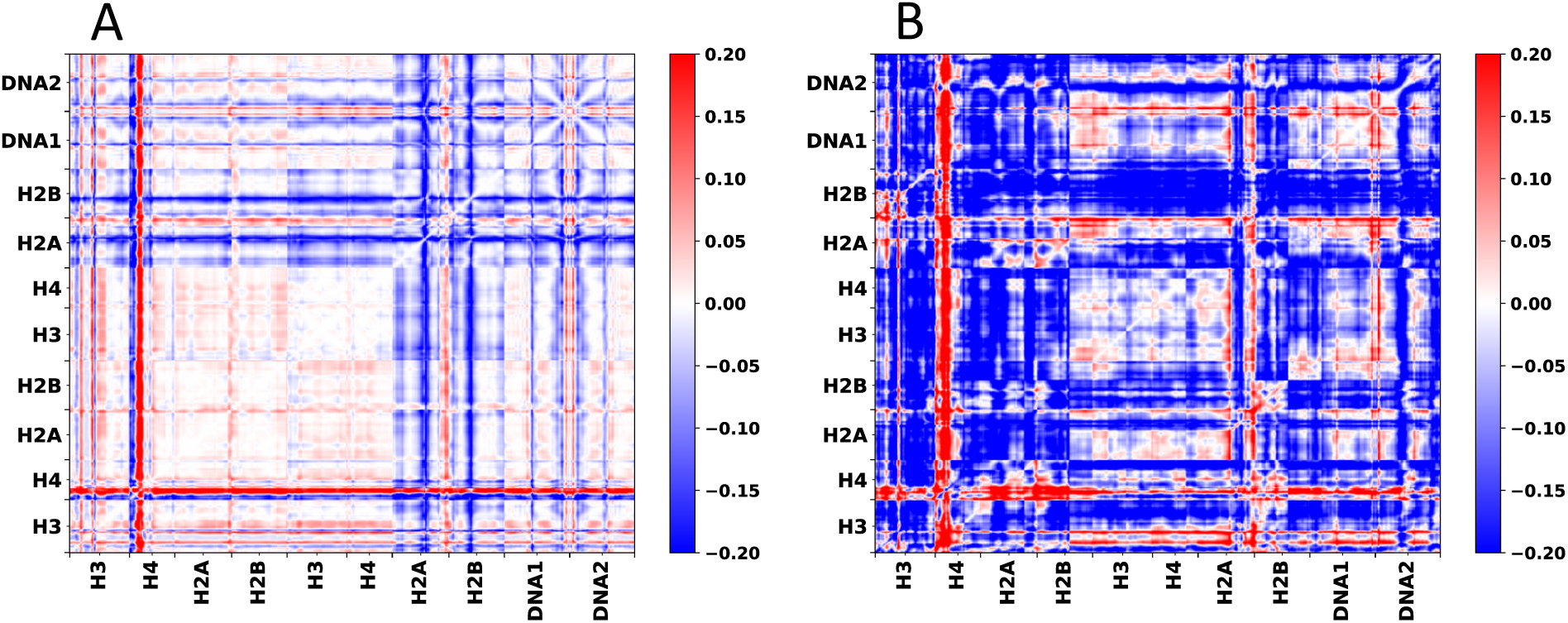
(A–B) Differences in atomic mutual information (dynamical correlations) between H2AK119ub and H2BK120ub nucleosomes, relative to the canonical system. Positive values reflect increased coupling; negative values reflect reduced coupling. Diagonal blocks represent intra-molecular correlations; off-diagonal blocks indicate inter-molecular correlations between histones and DNA.

In general, the changes introduced in H2BK120ub correlations were larger in magnitude than those in H2AK119ub, with multiple correlation changes in the range of 0.15-0.20 compared to more typical values of 0.05 in H2AK119ub. Together, these correlation analyses show that ubiquitination had a site-dependent effect on modulating the inter- and intra-nucleosome dynamics, with H2BK120ub producing a larger magnitude effect than H2AK119ub.

### Conformational Landscapes and Assembly Pathways of the Canonical Nucleosome

All-atom simulations revealed that both ubiquitinations influenced nucleosome structures and dynamics on the microsecond timescale. H2AK119ub stabilized nucleosome dynamics by promoting direct ubiquitin–DNA interactions and strengthening dimer association within the histone core, whereas H2BK120ub reduced dimer stability and increased overall nucleosome dynamics. We hypothesized that the differences in nucleosome dynamics observed across systems in all-atom simulations could drive large-scale structural variations such as nucleosome folding over longer timescales, which were not accessible to all-atom simulations. To test this, we performed coarse-grained simulations of canonical, H2AK119ub, and H2BK120ub nucleosomes. These simulations used the 3SPN.2C^52^ DNA and AICG2+ structure-based protein potentials, with the ubiquitin linkage parameterized by iterative Boltzmann inversion calculations (Figure S2), following the work of Terakawa and Takada.^56^ For each system, we performed 800 simulations of 10^8^ steps from nucleosomal, hexasomal, and tetrasomal states, and integrated the results into a Markov State Model (MSM) (see Methods for more details).

The MSM for the canonical NCP was clustered into 11 macrostates. For clarity, Figure 7 shows the six highest-flux pathways from tetrasomal to nucleosomal states, while the full network is shown in Figure S9. We identified three dominant tetrasomal states (T, T1, T2) that differed in DNA extension and dimer positioning. State T features fully extended DNA arms with one dimer at the end and the other closer to the tetramer. T1 is the most compact, with dimers nestled between the two DNA strands. T2 is intermediate, with both DNA arms and dimers nearer the core, and the DNA bent around one of the dimers. We also identified four major hexasomal states (HR, HR2, HR3, HL). In HR, the dimer is close to the core and unwrapped AT-poor DNA entry; HR1 positions the dimer at the DNA end; HL features unwrapping of the AT-rich DNA exit and dimer; and HR2/HR3 have intermediate conformations. The dominant nucleosome state is denoted N.

**Figure 7:**
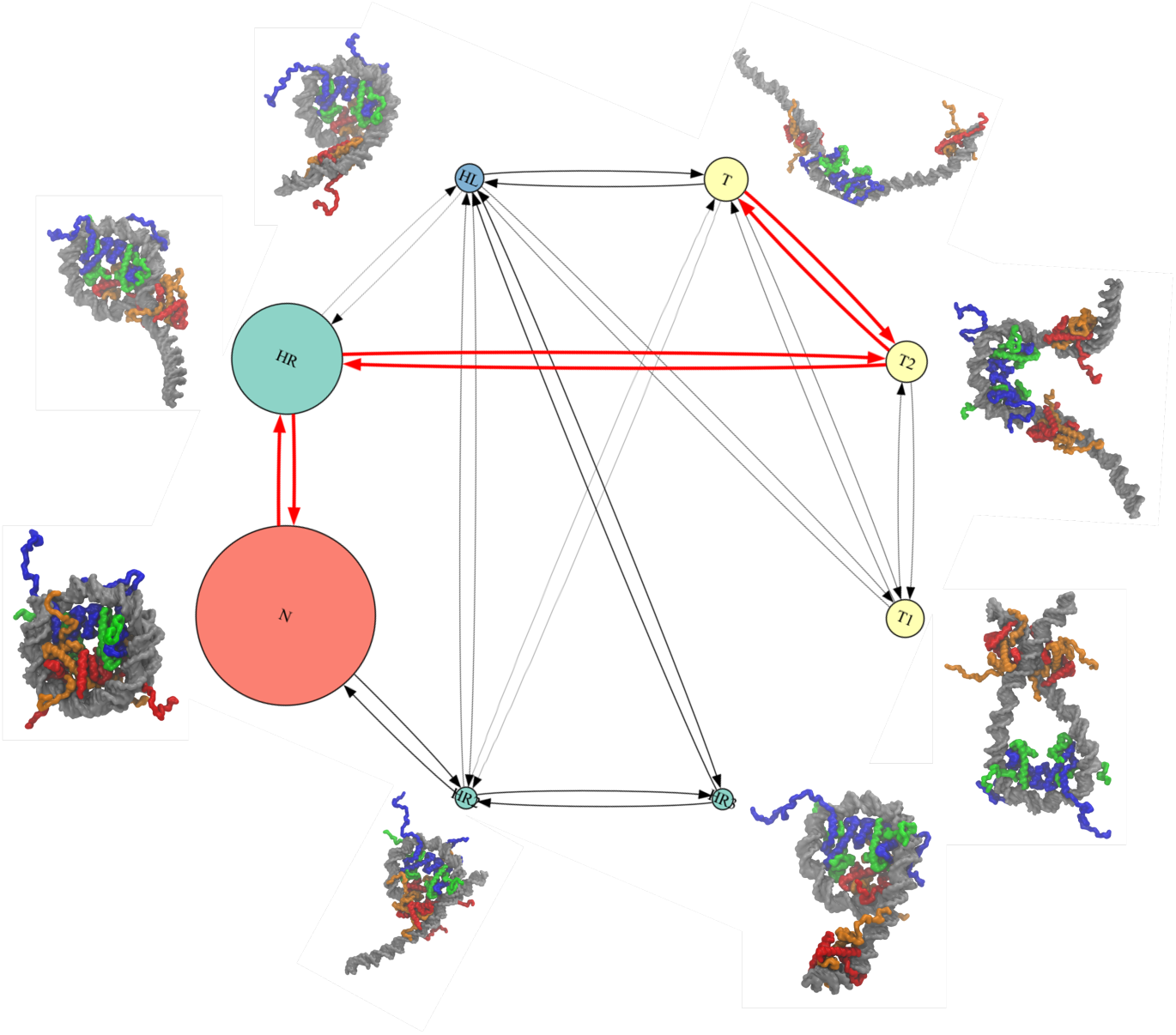
Top six assembly pathways for the canonical nucleosome, as identified by transition path theory from the Markov State Model. Each node represents a macrostate, with size proportional to equilibrium free energy. Arrow thickness indicates pathway dominance. State energies relative to the most probable state are listed in the accompanying table.

**Figure 8:**
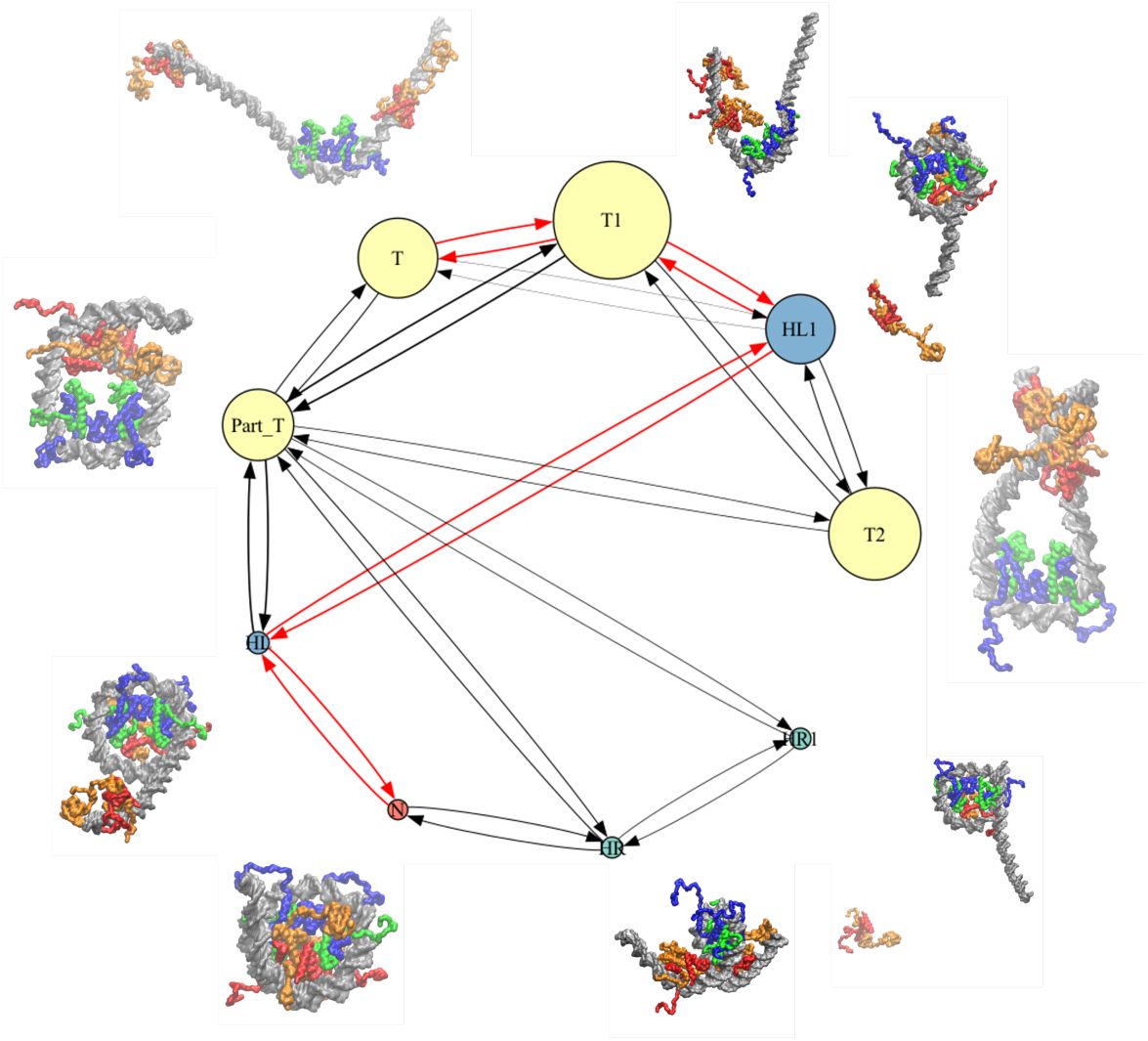
Top six assembly pathways for the H2AK119ub-modified nucleosome, as identified by transition path theory from Markov State Model analysis. Each node represents a macrostate, with its size proportional to the equilibrium free energy. The arrow thickness indicates the pathway’s dominance. State free energies relative to the most probable state are listed in the accompanying table.

Analysis of state free energies showed that the nucleosomal state is the global minimum, with the highest population. However, other macrostates also have relatively low free energies and are marginally populated at room temperature. For example, the HR hexasomal state is only 0.25*·*kcal/mol above the nucleosomal state, while HR2, HR3, and HL are 2.18–3.05*·*kcal/mol higher. This points to HR as the dominant hexasomal conformation. All three tetrasomal states have nearly identical free energies of 1.29–1.57*·*kcal/mol above the nucleosomal state, despite their differences in compactness. Together, these results suggest that the canonical coarse-grained system can exist in a population of states with varying degrees of compactness and DNA wrapping.

Pathway analysis revealed that folding from tetrasomal to nucleosomal states is dominated by a few key routes. The main pathway, T*↓*T2*↓*HR*↓*N includes folding of the open tetrasome to compact T2, transition to the low-free energy HR hexasome, and then nucleosome formation. This pathway accounts for 51.6% of the total flux between the tetrasome and nucleosome, and has a mean first passage time (MFPT) of 3.83*·*10^9^ MD steps (Table 3). The second dominant pathway bypasses HR and proceeds through HL, HR3, HR2, and then N. However, it is much slower with an MFPT an order of magnitude higher than the first pathway (1.19*·*10^10^) and represents 16.6% of the total flux. A third pathway resembles the main route but passes through T1 before reaching T2, increasing the MFPT to 2.09*·*10^10^ steps and representing 9.5% of the total flux. Notably, the enrichment of HR states along the folding pathways, which is driven by DNA entry-side unwrapping, matches the increased entry-side breathing seen in all-atom simulations.

**Table 3:** Dominant folding pathways and mean first passage times (MFPT) for assembly from tetrasome to nucleosome in the canonical system, determined by Markov State Model analysis. The percentage indicates the fraction of total folding flux through each pathway.

| Rank | Pathway | Mean First Passage Time | % of Total Flux |
| --- | --- | --- | --- |
| 1 | T→T2→HR→N | $3.83 \cdot 10^9$ | 51.6% |
| 2 | T→HL→HR3→HR2→N | $1.19 \cdot 10^{10}$ | 16.6% |
| 3 | T→T1→T2→HR→N | $2.09 \cdot 10^{10}$ | 9.5% |
| 4 | T→T1→HL→HR1→N | $2.09 \cdot 10^{10}$ | 9.4% |
| 5 | T→HR1→N | $5.31 \cdot 10^{10}$ | 3.7% |
| 6 | T→T1→HL→HR→N | $5.37 \cdot 10^{10}$ | 2.1% |

**Table 4:** Dominant folding pathways and mean first passage times (MFPT) for assembly from tetrasome to nucleosome in the H2AK119ub system, determined by Markov State Model analysis. The percentage indicates the fraction of total folding flux through each pathway.

| Rank | Pathway | Mean First Passage Time | % of Total Flux |
| --- | --- | --- | --- |
| 1 | T→T1→HL1→HL→N | $8.86 \cdot 10^{10}$ | 24.7% |
| 2 | T→T1→Part_T→HL→N | $9.00 \cdot 10^{10}$ | 24.3% |
| 3 | T→T1→T2→Part_T→HL1→HL→N | $1.37 \cdot 10^{11}$ | 16.0% |
| 4 | T→Part_T→HR→N | $1.47 \cdot 10^{11}$ | 15% |
| 5 | T→T1→T2→Part_T→HR1→HR→N | $1.87 \cdot 10^{11}$ | 11.7% |
| 6 | T→HL1→HL→N | $4.86 \cdot 10^{11}$ | 4.5% |

Overall, the folding landscape of the canonical nucleosome is dominated by sequential compaction of the core and a preference for hexasome intermediates where the DNA exit strand is folded, with most flux passing through a small set of low-energy pathways.

### Site-Specific Ubiquitination Destabilizes Nucleosomes and Slows Folding

To understand how ubiquitination affects large-scale nucleosome structures, we performed coarse-grained simulations and constructed Markov State Models for H2AK119 and H2BK120 ubiquinated nucleosomes. In general, the conformations of macrostates identified in these analyses were similar to those in the canonical system. However, ubiquitination significantly altered the free energies of these states and the transitions between them.

In the H2AK119-ubiquitinated system, the nucleosomal state was significantly destabilized compared to the tetrasomal states. The extended tetrasomal states T and T1 had free energies of 0.00 and 0.64 kcal/mol, while the more compact tetrasomal states T2 and Part T had similar low free energies of 0.36 and 0.87 kcal/mol (Table 5). In contrast, the nucleosomal state had a much higher free energy of 6.54 kcal/mol. Most hexasomal states had free energies between 5.70 and 6.12 kcal/mol, with the exception of HL1, which was significantly lower at 0.95 kcal/mol.

**Table 5:** Relative free energies (kcal/mol) of principal macrostates along the nucleosome folding pathway in the H2AK119ub system, calculated from Markov State Model analysis. Energies are given relative to the most stable state.

| State | Energy (Kcal/mol) |
| --- | --- |
| Tetrasome (Extended) | T (0.64), T1 (0.00), |
| Tetrasome (Compact) | T2 (0.36), Part_T (0.87) |
| Hexasome Right (HR) -<br>DNA-exit-side folded | HR (6.12), HR1 (6.11) |
| Hexasome Left (HL) -<br>DNA-entry-side folded | HL (5.70), HL1 (0.95) |
| Nucleosome (N) | N (6.54) |

**Table 6:** Dominant folding pathways and mean first passage times (MFPT) for assembly from tetrasome to nucleosome in the H2BK120ub system, determined by Markov State Model analysis. The percentage indicates the fraction of total folding flux through each pathway.

| Rank | Pathway | Mean First Passage Time | % of Total Flux |
| --- | --- | --- | --- |
| 1 | T→T1→HL→N | $5.03 \cdot 10^{11}$ | 0.3% |
| 2 | T→Part_T→HL→N | $8.61 \cdot 10^{11}$ | 0.2% |
| 3 | T→Part_T→T1→HL→N | $9.80 \cdot 10^{11}$ | 0.2% |
| 4 | T→HR→N | $1.58 \cdot 10^{12}$ | 0.1% |
| 5 | T→T2→T5→T4→HL→N | $2.32 \cdot 10^{12}$ | 0.1% |
| 6 | T→T3→HR→HL→N | $2.55 \cdot 10^{12}$ | 0.1% |

The folding pathways for the H2AK119ub system were slower than in the canonical system. While there were more folding pathways with appreciable flux, only a handful contributed the majority of transitions between the tetrasomal and nucleosomal states. The two fastest folding pathways had nearly identical MFPTs of around 9.0*·*10^10^ timesteps, each representing 24.7% and 24.3% of the total flux, respectively. The first pathway transitioned from the extended T to T1 tetrasomal states, then folded into the left hexasome states HL1 and H1 before reaching the N nucleosome state. The second pathway was similar, however, instead of folding through HL1, it passed through the tetrasomal Part T state. Pathways three through five contributed significant flux (11.7%-16.0%) with MFPT ranging from 1.37*·*10^11^ to 1.87*·*10^11^ timesteps.

Ubiquitination at H2BK120 resulted in even greater destabilization of the nucleosomal state. In the H2BK120ub MSM the nucleosome was sparsely populated, with a free energy of 10.01 kcal/mol. Multiple extended and compact tetrasomal states had low free energies ranging from 0.00 to 1.49 kcal/mol. Two hexasomal states were observed, one with the hexamer on the entry DNA and the other on the exit DNA. Both ad free energies of 3.28 kcal/mol, intermediate between the tetramer and nucleosome states (Table 7).

**Table 7:** Relative free energies (kcal/mol) of principal macrostates along the nucleosome folding pathway in the H2BK120ub system, calculated from Markov State Model analysis. Energies are given relative to the most stable state.

| State | Energy (Kcal/mol) |
| --- | --- |
| Tetrasome (Extended) | T (1.10), T2 (0.51), T5(0.00) |
| Tetrasome (Compact) | Part_T(1.13), T1 (0.78), T3 (1.49), T4 (1.13) |
| Hexasome Right (HR) - DNA-exit-side folded | HR (3.28) |
| Hexasome Left (HL) - DNA-entry-side folded | HL (3.28) |
| Nucleosome (N) | N (10.01) |

The nucleosome assembly pathway in H2BK120ub was highly diffuse, with many interconnected states rather than a small number of dominant routes (Figures 9 and S11). The folding times were also much slower: the fastest MFPT was 5.03*·*10^11^ steps, two orders of magnitude greater than the canonical and one order greater than the H2AK119ub system (Table 6). Although both the HL and HR states had similar free energies, the top three dominant pathways all transitioned from the tetrasomal to the nucleosomal state through HL. The dominant pathway, however, accounted for only 0.3% of the total flux, indicating that nucleosome folding in H2BK120ub is highly heterogeneous and occurs much less efficiently than in the canonical or H2AK119ub systems.

**Figure 9:**
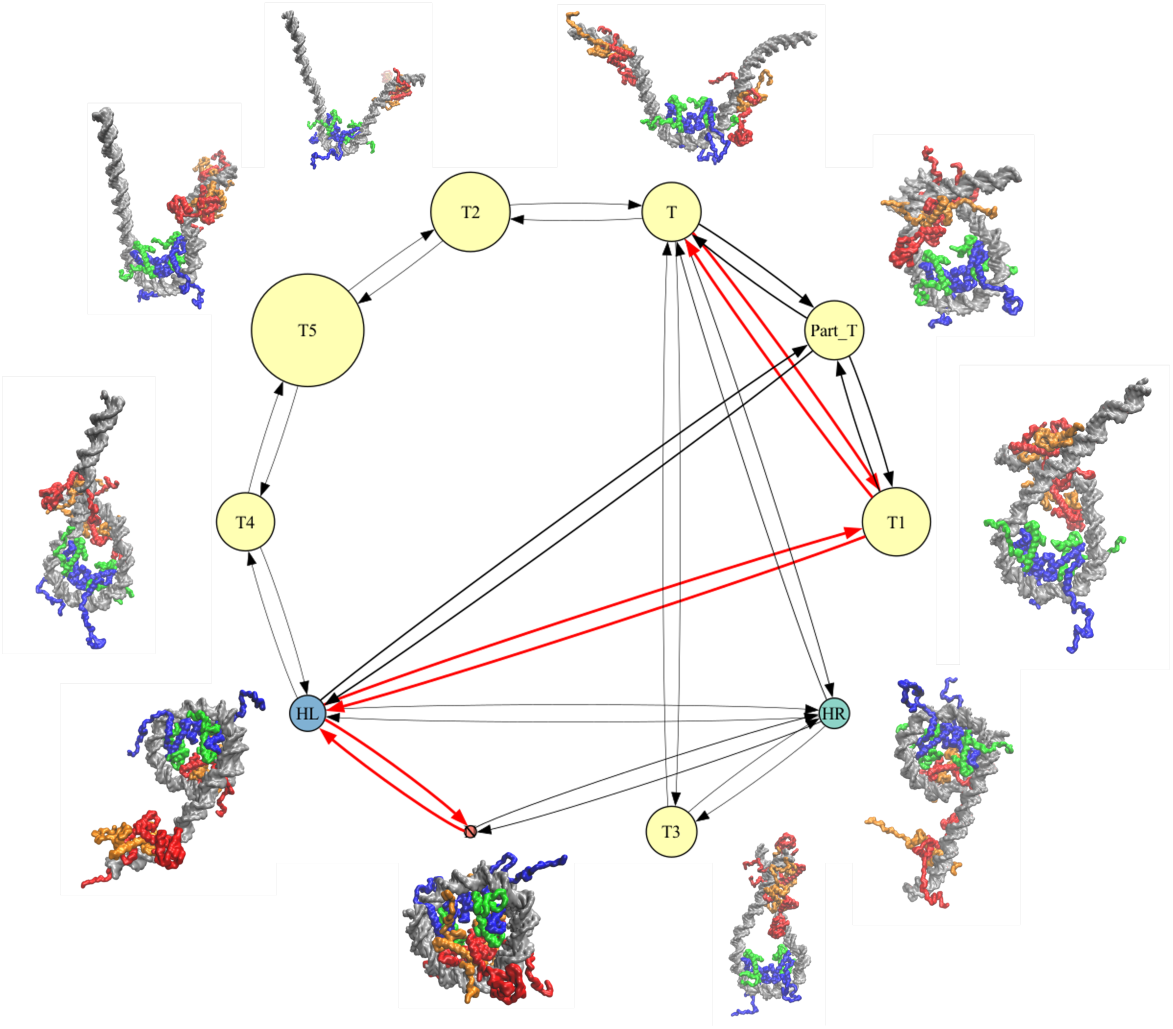
Top six assembly pathways for the H2BK120ub-modified nucleosome, as identified by transition path theory from Markov State Model analysis. Each node represents a macrostate, with its size proportional to the equilibrium free energy. The arrow thickness indicates the pathway’s dominance. State free energies relative to the most probable state are listed in the accompanying table.

**Figure 10:**
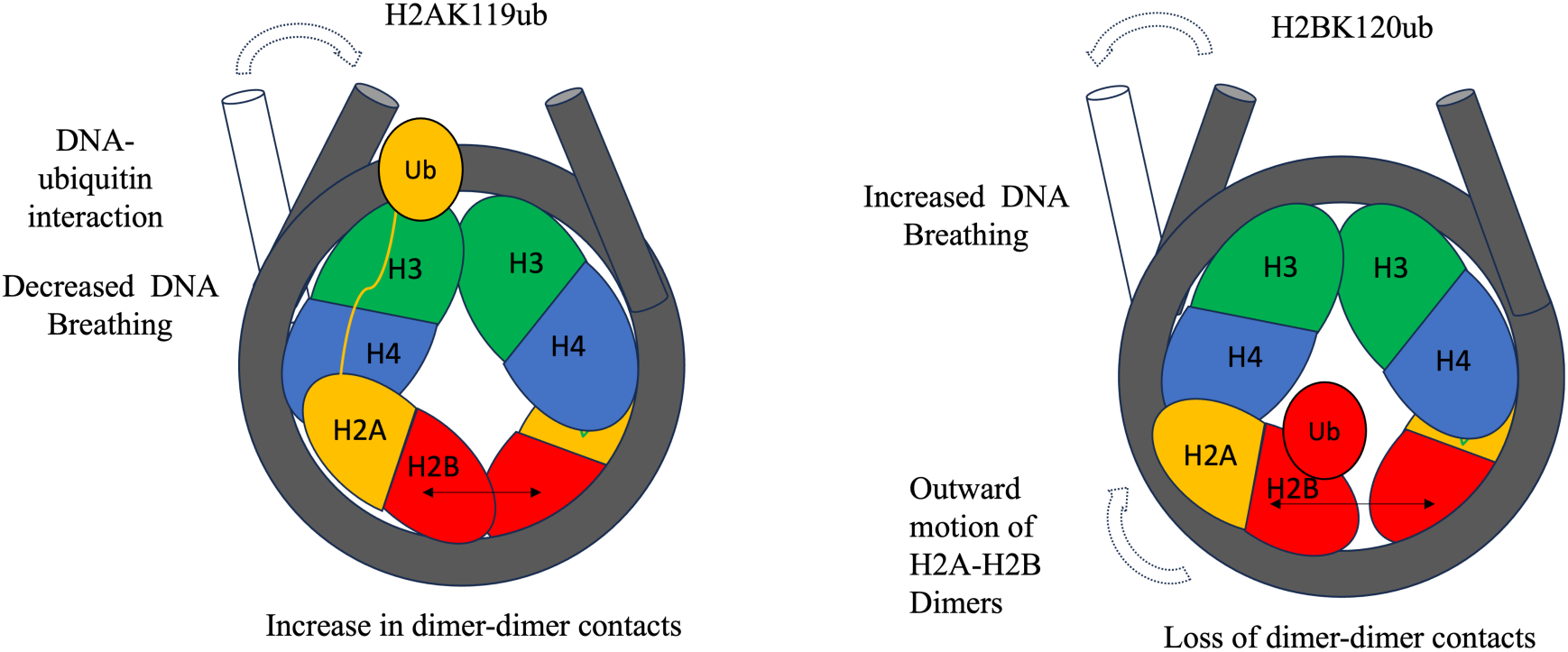
Illustration depicting the distinct mechanisms by which H2AK119ub and H2BK120ub alter nucleosome stability and assembly. H2AK119ub indirectly strengthens the L1–L1 interface between H2A histones, stabilizing the nucleosome core and slowing assembly. In contrast, H2BK120ub disrupts these contacts and weakens the histone core, promoting partially assembled hexasome and tetrasome states. Arrows and annotations indicate the sites of ubiquitination, key affected interfaces, and the resulting structural and kinetic effects on nucleosome folding.

## Discussion

Ubiquitination of histones at specific sites, such as H2AK119 and H2BK120, is associated with transcriptionally inactive or active chromatin states, respectively^23,26^, respectively. Experimental studies have shown that ubiquitin directly impacts nucleosome stability^69^, but the molecular basis for these effects remains elusive. In this study, we employed both all-atom and coarse-grained simulations to probe how site-specific ubiquitination alters nucleosome dynamics across microsecond to second timescales, which are periods directly relevant to nucleosome assembly and stability.

Our results indicate that ubiquitination alters the motions of H2A/H2B dimers within the nucleosome core in a site-specific manner. H2AK119ub interacts favorably with the DNA entry and exit regions, whereas ubiquitin at H2BK120 remains close to the histone core with minimal DNA interactions, as demonstrated in a recent cryo-EM study^70^. In the H2BK120ub system, both dimers undergo pronounced outward tilting relative to the dyad axis. In the H2AK119ub system, this effect is asymmetric: one dimer shows behavior similar to the canonical nucleosome, while the other exhibits increased tilting angles over 2 µs of simulation (Figure 3).

Ubiquitination profoundly impacts the H2A L1-L1 interface, a region directly implicated in overall nucleosomal stability.^71,72^ The H2A L1 loops are comprised of four residues located at the base of the nucleosome near SHLs *±*4, and are the sole points of contact between the two dimers. Changes in the L1-L1 interface have been shown to alter nucleosome stability, as we previously demonstrated in the macroH2A variant^72,73^. Our simulation show that H2BK120ub significantly disrupts L1-L1 contacts, while H2AK119ub reorganizes these dimer-dimer as well as other dimer-tetramer contacts (Figures S6-S7). Disruption of these contacts in H2BK120ub leads to decoupling of dimer, tetramer, and DNA motions (Figure 6), and in both systems this may contribute to the observed increase in dimer orientations relative to the canonical nucleosome.

Our coarse-grained simulations, which mirror experiments where H2A/H2B dimers are added to pre-formed tetrasomes^26,59^, show that ubiquitination does not alter the basic folding pathway from tetramer to hexasome to nucleosome. However, it drastically slows this process. H2AK119ub slows nucleosome folding by an order of magnitude (Table 4), while H2BK120ub produces a two-order-of-magnitude slowdown (Table 6). In both cases, ubiquitinated systems favor hexasomal and tetrasomal intermediates over fully formed nucleosomes (Tables 5 and 7). For example, in the canonical nucleosome, the most stable hexasomal state has a free energy similar to that of the nucleosome. In contrast, for H2AK119ub and H2BK120ub, the most stable hexasomal states are approximately 6 kcal/mol more favorable than the nucleosomal state. In both modified systems, the tetrasomal state is the most energetically favored overall. This substantial energy difference shifts the equilibrium toward tetrasomal and hexasomal states when the nucleosome is ubiquitinated.

Our finding that di-ubiquitination destabilizes the nucleosome and promotes more dynamic states is consistent with recent single-molecule experiments. Luo *et al.* showed with single-molecule magnetic tweezer experiments that monoubiquintated H2BK120 nucleosomes have lower mechanical stability than their canonical counterparts, which increases the affinity of the FACT complex and helps maintain the transcriptionally active state.^26^ They also reported that the free energy difference of unwrapping the outer fold of DNA was on the order of 5 kcal/mol, comparable to our MSM-predicted differences between hexasomal and tetrasomal states. Although this work focused on monoubiquitinated nucleosomes, one would expect di-ubiquitination to have similar or stronger destabilizing effects.

While our simulations capture both short and long timescales, the coarse-grained approach has several limitations that may impact accuracy. For example, it is a simplified model in which the ionic atmosphere is approximated using the Debye–Hückel formalism^74^. Secondary structure changes are not modeled, and native contacts are assumed, although some are disrupted in all-atom simulations over microsecond timescales. While these factors may affect the numerical precision of our MSMs, similar models have shown good agreement with experimental results^75,76^, supporting their use for uncovering biophysical mechanisms in comparable systems.

In summary, our study provides a mechanistic framework for how site-specific ubiquitination at H2AK119 and H2BK120 regulates nucleosome dynamics and stability, showing that these modifications are not just passive epigenetic marks but active determinants of chromatin physical properties. The observation that both modifications favor partially assembled nucleosome states suggests a direct route by which ubiquitin marks could modulate chromatin accessibility *in vivo*, with implications for transcription, replication, and repair. This prediction is readily testable by single-molecule FRET or force spectroscopy. By clarifying the physical impact of these distinct ubiquitin modifications, our work helps bridge the gap between structural biophysics and biological function, and sets the stage for future experimental validation in living cells.

## Acknowledgments

This project was supported by the National Institutes of Health grant R35GM119647.

## Data availability

All simulation inputs, scripts used for analysis, and processed data files are available at https://github.com/WereszczynskiGroup/supplemental-data-baweja-ubiquitin-2025. Raw trajectory data generated in this study are available on Zenodo at https://doi.org/10.5281/zenodo.14866729. These trajectories are water-stripped and temporally strided to reduce file size but cover the full duration of each simulation. Together, these resources are sufficient to reproduce all analyses and key results reported in the manuscript.

