## Supporting Information for "Mechanistic basis for the opposing effects of H2A and H2B ubiquitination on nucleosome stability and dynamics"

Table S1: All-atom Simulation Details

| **System** | **Number of Simulations** | **Simulation Time (μs)** | **Total Sampling (μs)** |
|---|---|---|---|
| Canonical | 3 | 2.0 | 6.0 |
| H2AK119ub | 3 | 2.0 | 6.0 |
| H2BK120ub | 3 | 2.0 | 6.0 |

Table S2: Coarse-grain Simulation Details

| **System** | **Number of Simulations** | **Time steps** |
|---|---|---|
| Canonical | 800 | $10^8$ |
| H2AK119ub | 800 | $10^8$ |
| H2BK120ub | 800 | $10^8$ |

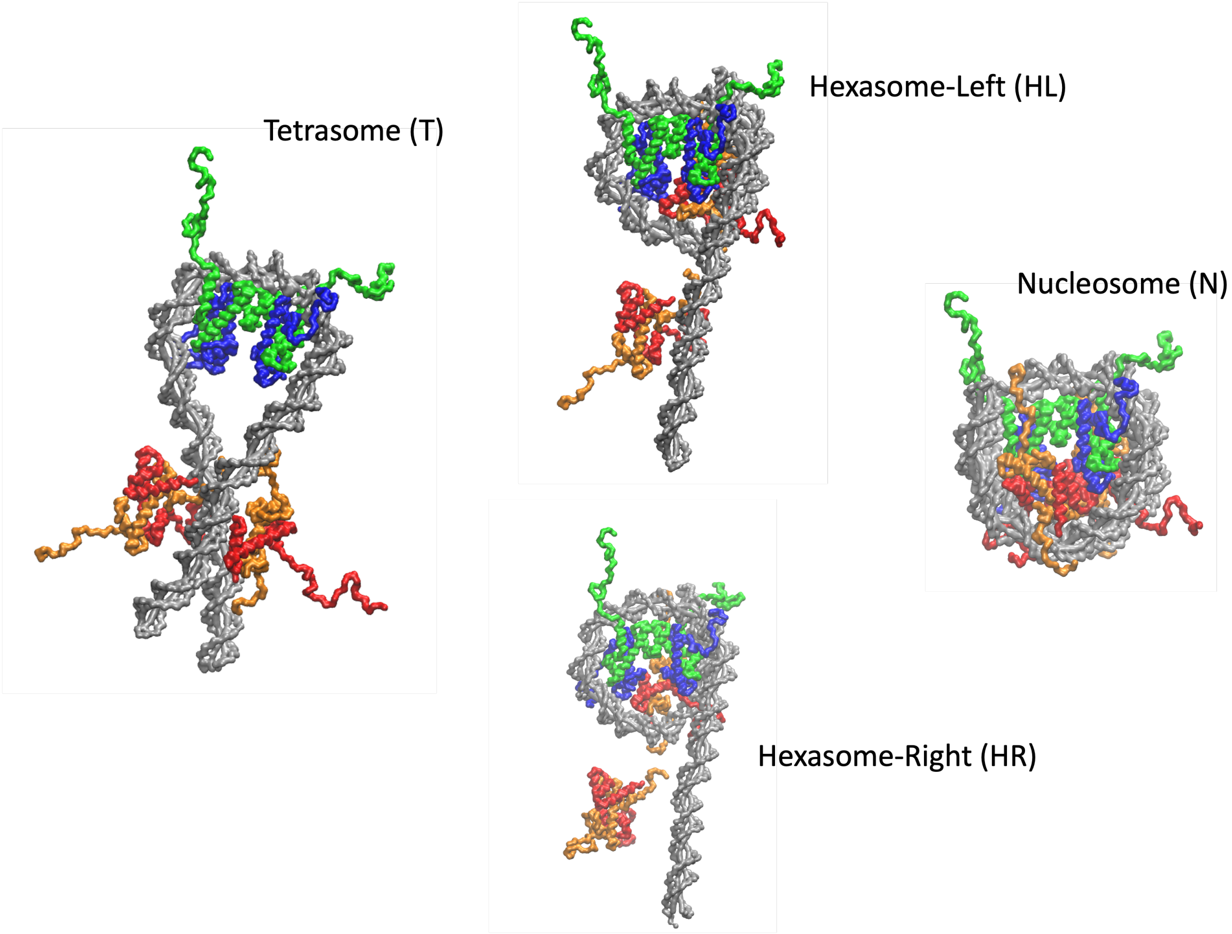


Figure S1: Initial Conformations of the Tetrasome (T), Hexasome-Left (HL), Hexasome-Right (HR) and Nucleosome (N)

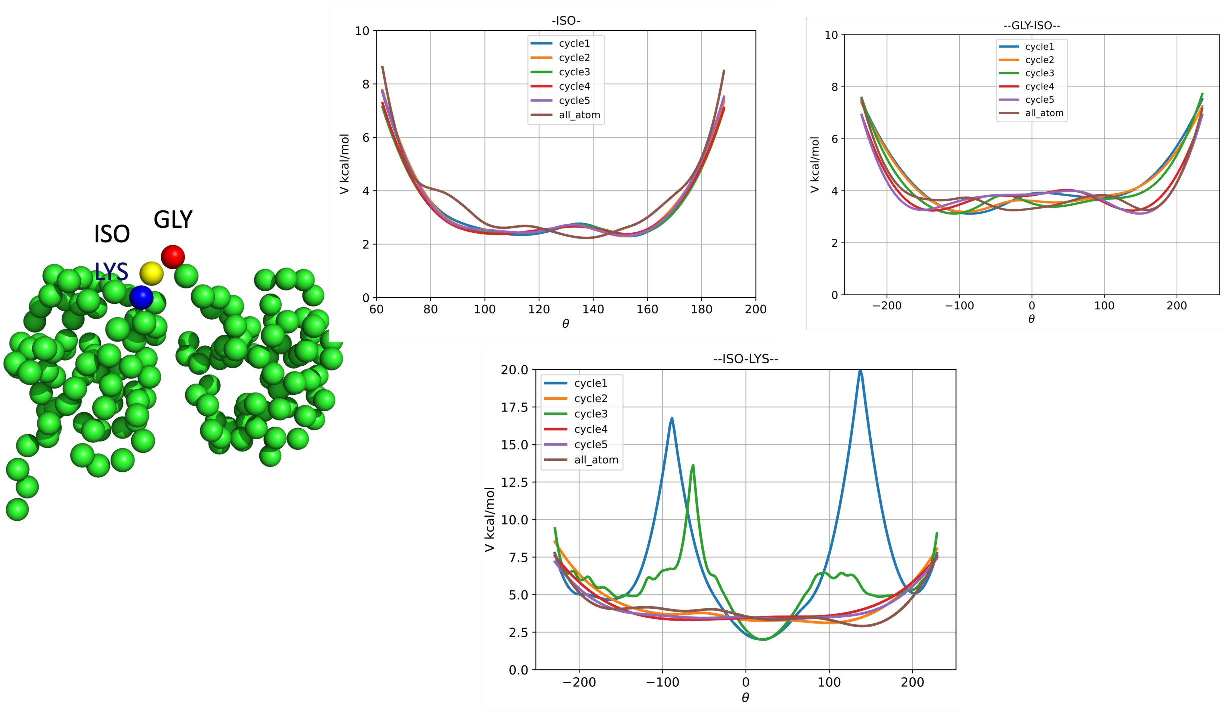


Figure S2: Bond and dihedral angles distribution obtained from Boltzmann Inversion protocol

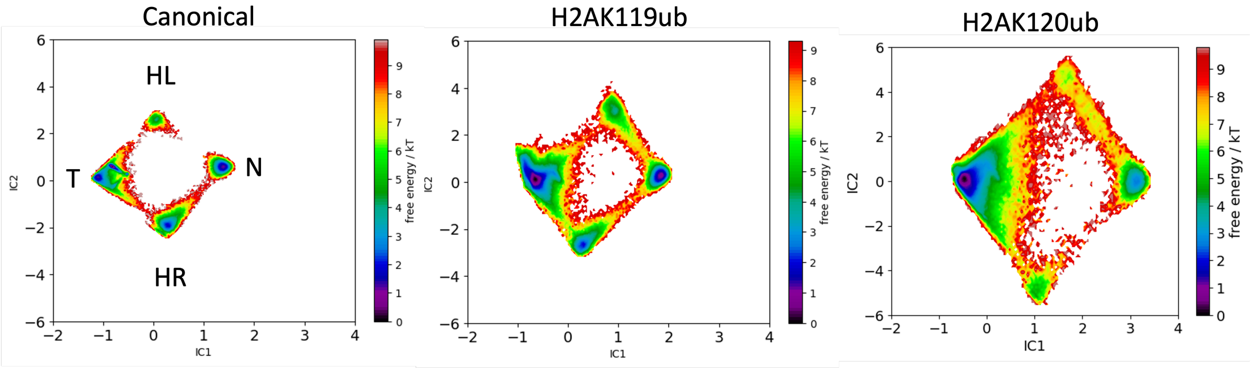


Figure S3: Projection along first two TICA coordinates revealed 4 metastable states tetrasome, hexasome-left, hexasome-right, and nucleosome

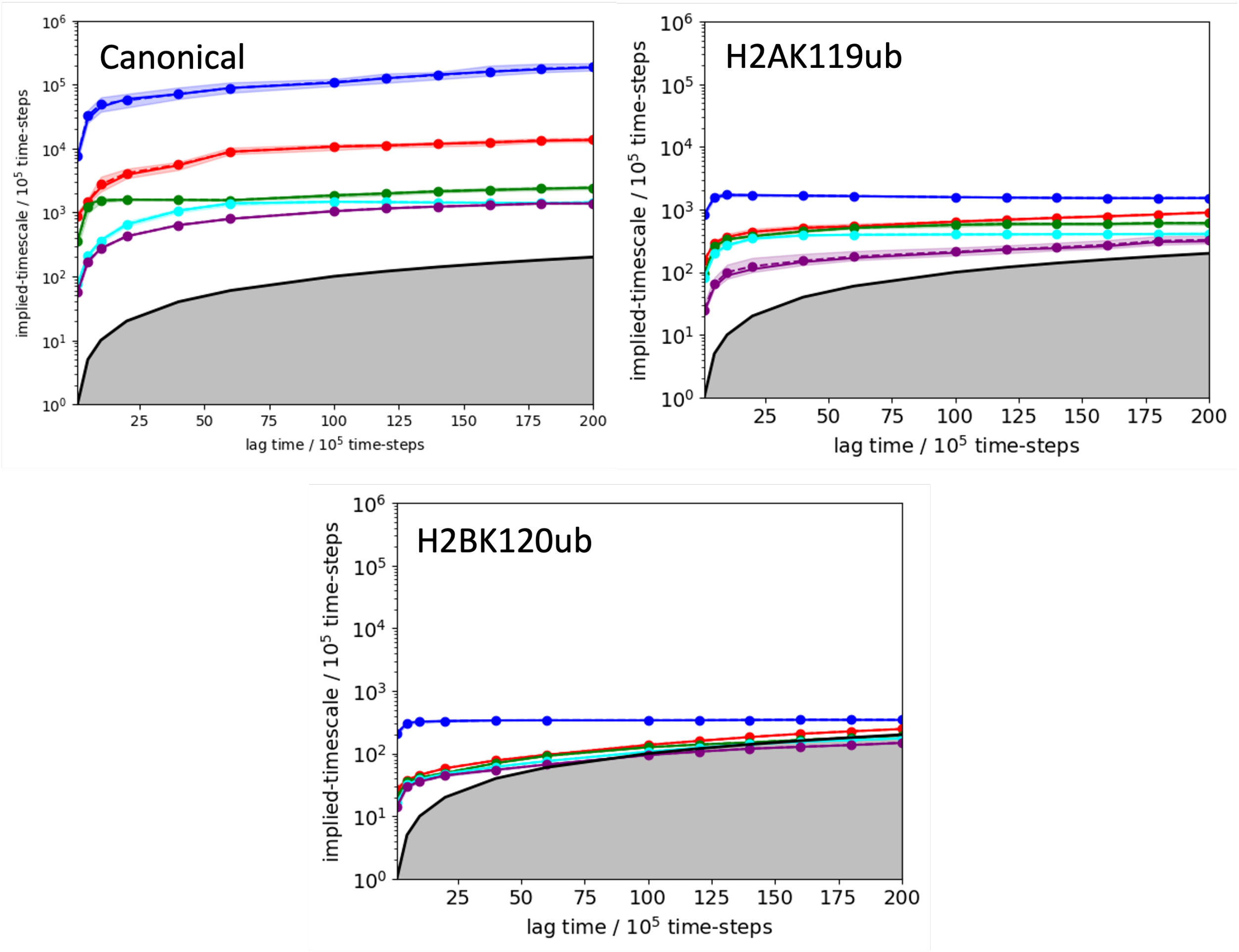


Figure S4: Implied time scales of the Markov State Model (MSM) as a function of the chosen lag-time. For the final model, we use a lag-time of $2 \times 10^6$ MD steps, after which most of the implied time scales have essentially converged. The shaded regions indicate 95% confidence intervals

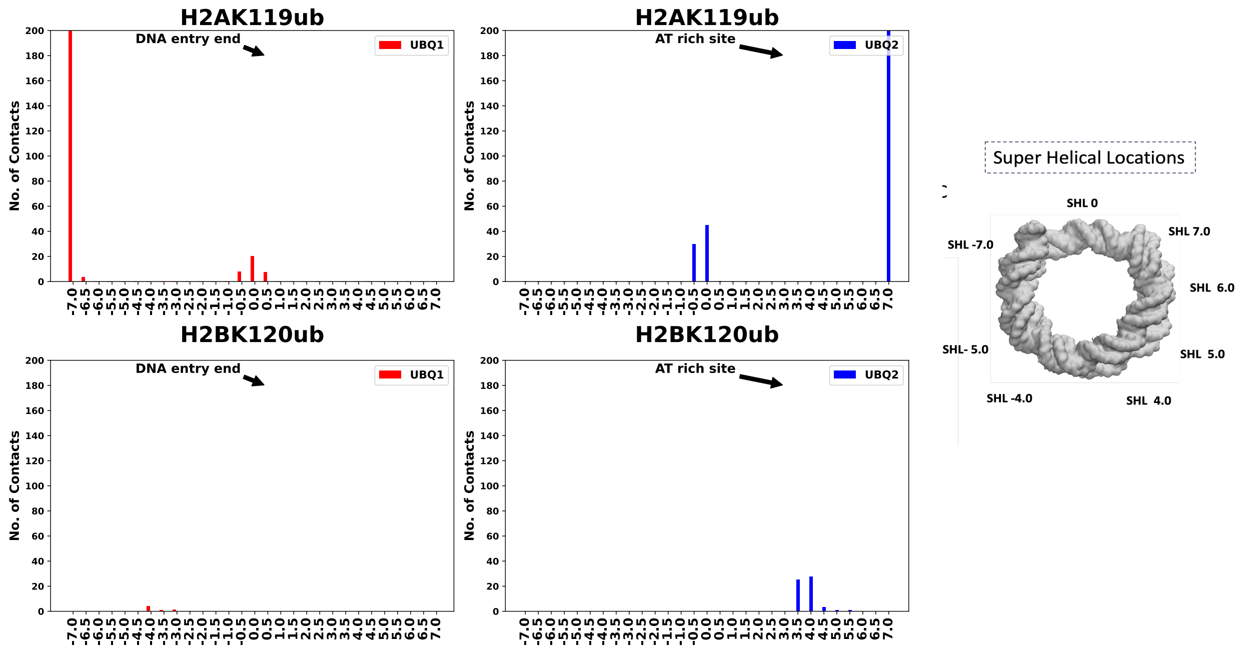


Figure S5: Interactions of ubiquitins with super helical locations in different systems. a) H2AK119 b) H2BK120

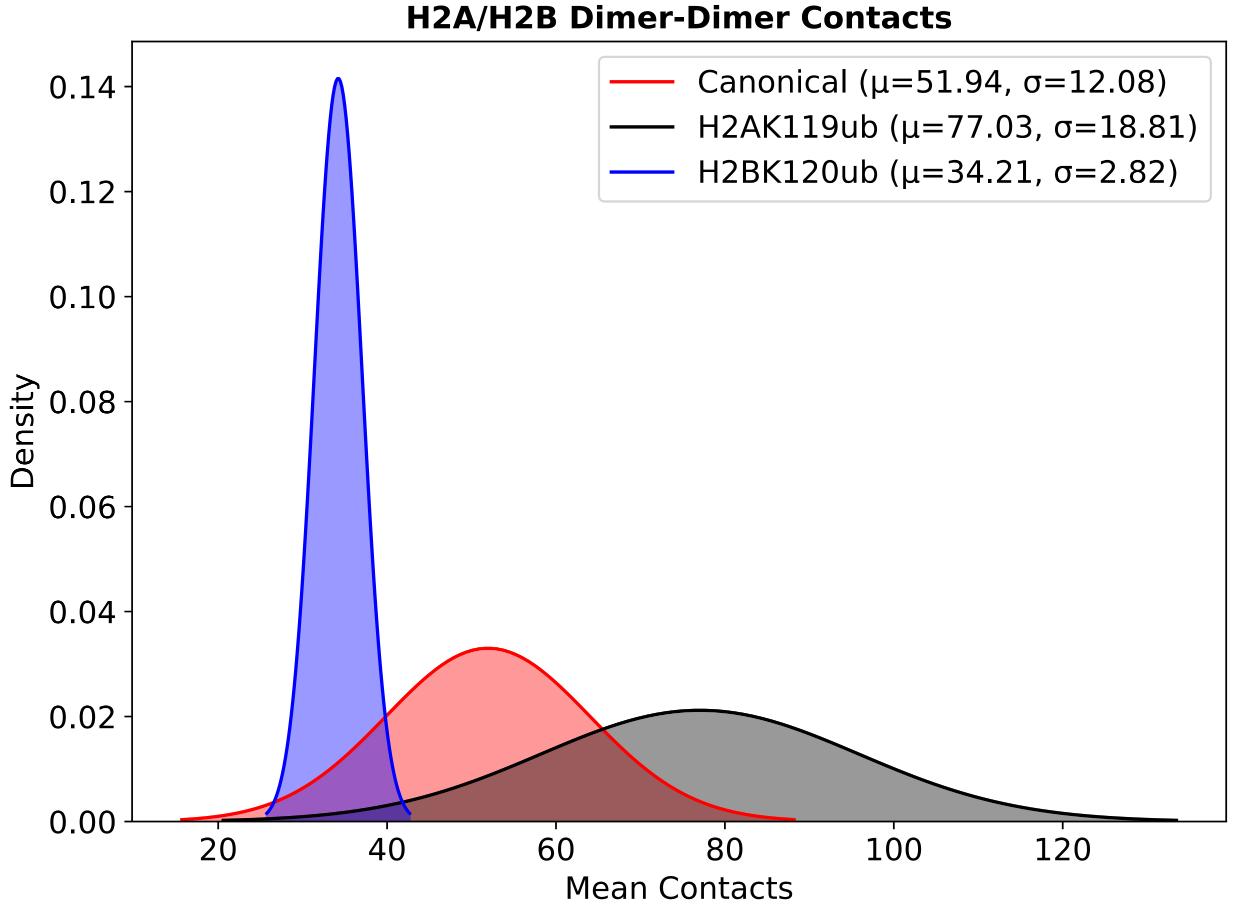


Figure S6: Mean number of dimer-dimer contacts of all the systems obtained by averaging the last 1.5 μs of three independent trajectories

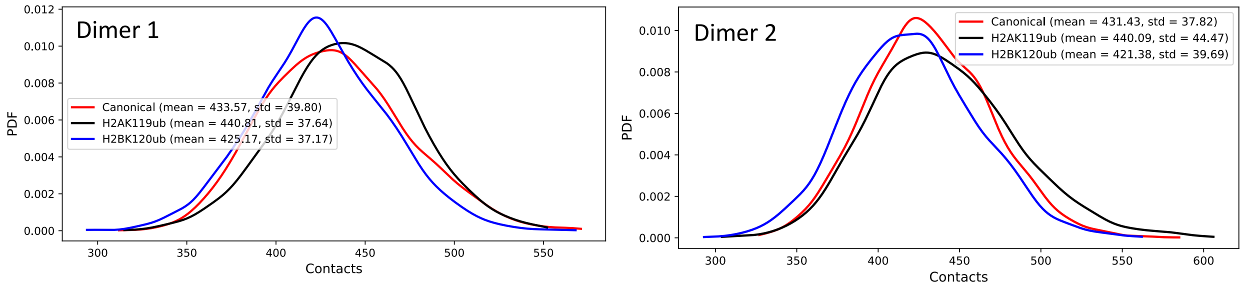


Figure S7: Distribution of mean number of dimer-tetramer contacts of all the systems obtained by averaging the last 1.5 μs of three independent trajectories

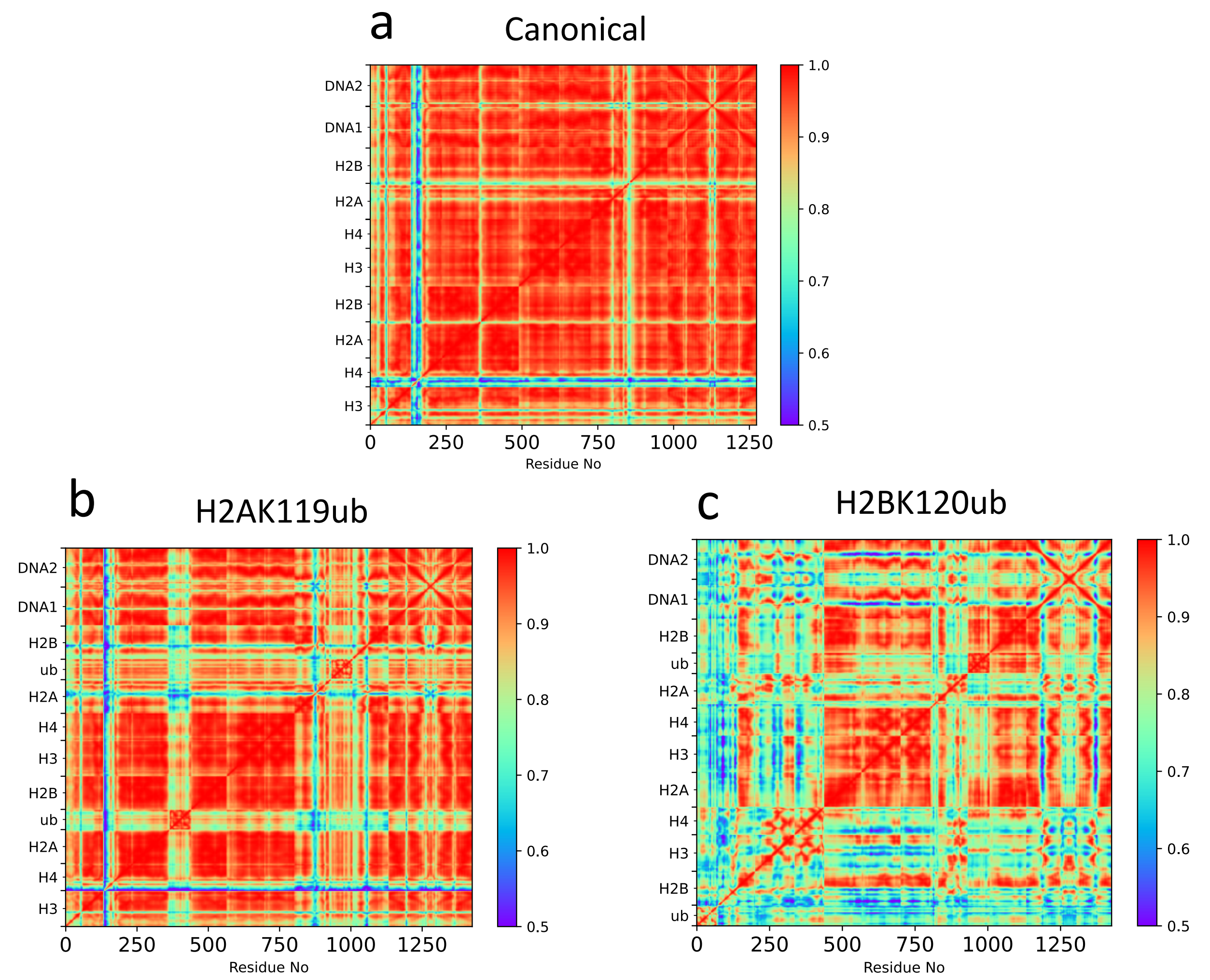


Figure S8: Mutual information of different systems a) Canonical, b) H2AK119ub, c) H2BK120ub

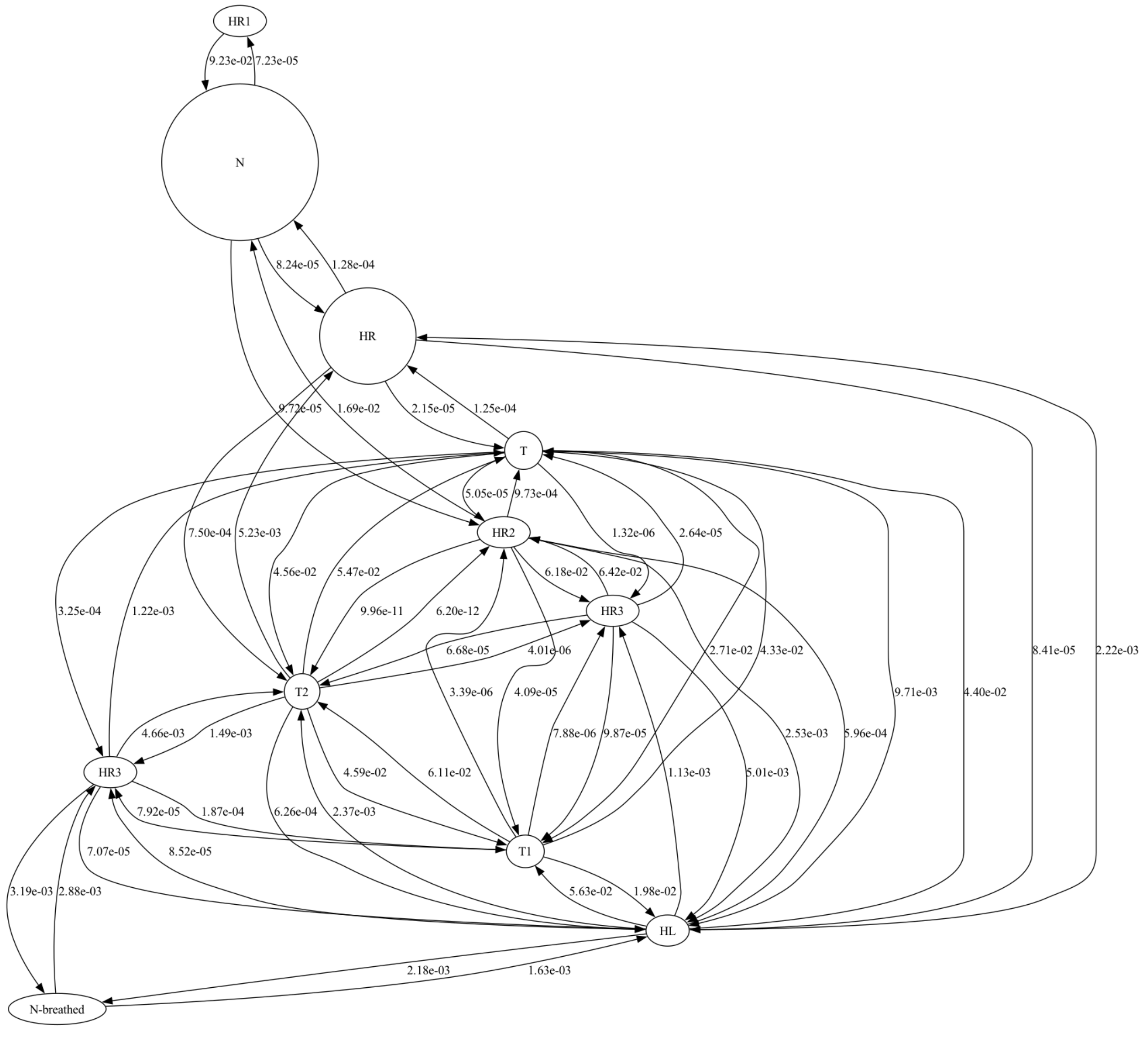


Figure S9: Markov state model of canonical system with transition pathways with transition probabilities between states

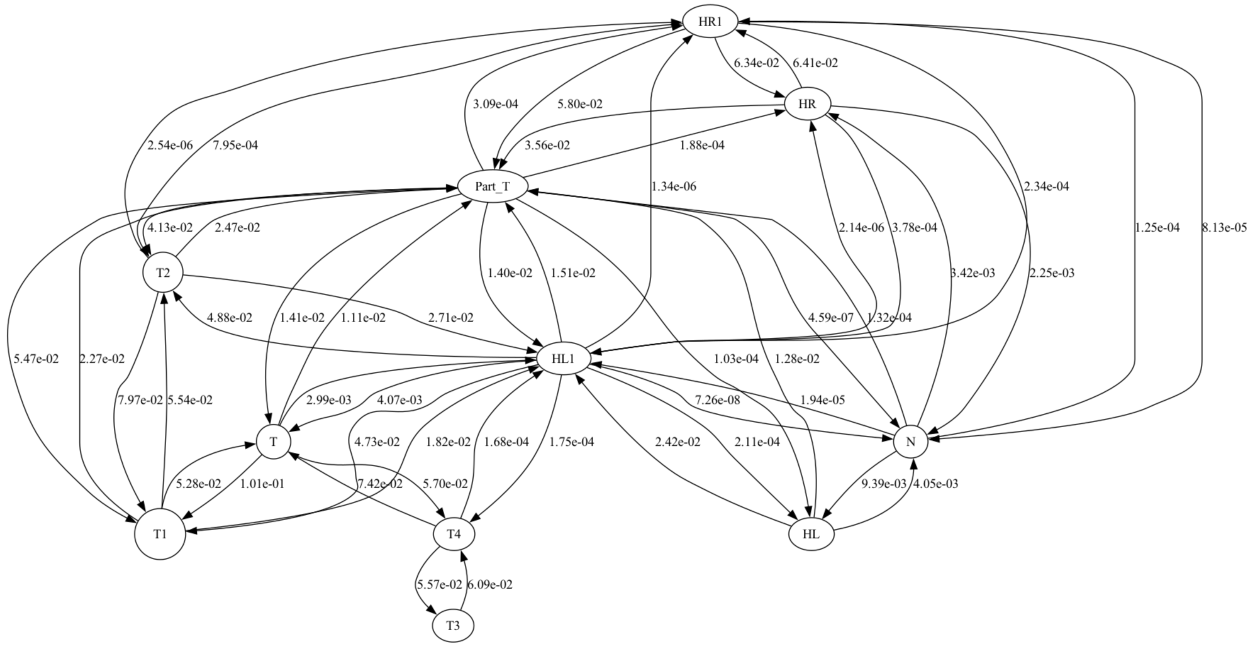


Figure S10: Markov state model of H2A119 system with transition pathways with transition probabilities between states

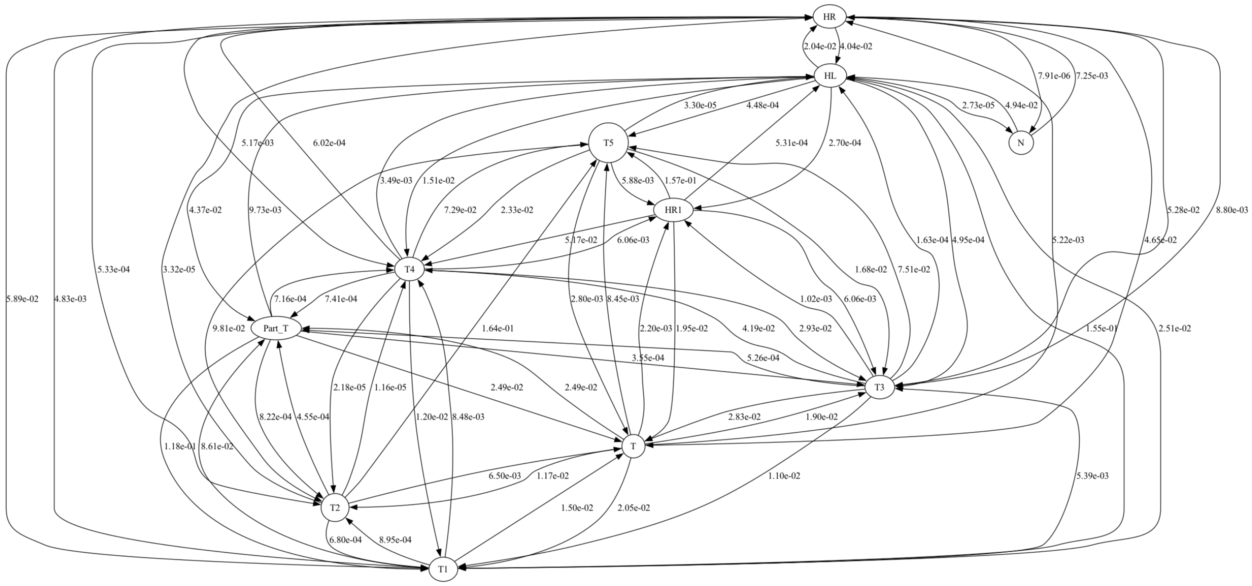


Figure S11: Markov state model of H2BK120 system with transition pathways with transition probabilities between states
